# *MIR192* Upregulates GLP-1 Receptor and Improves Statin-Induced Impairment of Insulin Secretion

**DOI:** 10.1101/2025.03.18.643960

**Authors:** Yu-Lin Kuang, Cassandra A.A. Locatelli, Yuanyuan Qin, Yuqing Zhang, Elizabeth Theusch, Antonio Muñoz-Howell, Gabriela Sanchez, Meng Lu, My-Anh Nguyen, Tanvi Yalamanchili, Xuanwen Wang, Gilbert Nalula, Aras N. Mattis, Akinyemi Oni-Orisan, Carlos Iribarren, Ronald M. Krauss, Erin E. Mulvihill, Marisa W. Medina

## Abstract

Statins are a commonly prescribed cholesterol lowering drug class that can increase the risk of new-onset diabetes (NOD). To investigate the molecular mechanisms underlying this effect, we generated human induced pluripotent stem cells (iPSCs) from individuals identified from electronic health records of Kaiser Permanente of Northern California who were susceptible to developing NOD after statin initiation or controls who maintained stable fasting glucose on statin treatment. RNA-seq analysis of iPSCs incubated with atorvastatin, simvastatin or mock buffer for 24 hours identified the long non-coding RNA *MIR194-2HG* as a top candidate gene. Statin-induced increases in its expression were observed in NOD resistant controls, while statin-induced reductions occurred in NOD susceptible cases. *MIR194-2HG* encompasses two microRNA genes: *MIR192* and *MIR194-2*. The mature microRNA miR-192-5p, derived from the 5’ arm of *MIR192*, was predicted to bind the 3’UTR of the glucagon like peptide 1 (GLP-1) receptor (*GLP1R*) transcript. Transfection of a rat insulinoma cell line INS-1 with a miR-192-5p mimic increased *Glp1r* transcript (1.41-fold) and protein (1.51-fold) levels compared to a scrambled control. Using a luciferase reporter containing the human *GLP1R* 3’UTR, miR-192-5p overexpression similarly increased luciferase signal (1.44-fold). The miR-192-5p mimic enhanced glucose stimulated insulin secretion (GSIS) in response to GLP1R agonists (1.64-1.81-fold) and rescued simvastatin-induced GSIS impairment in INS-1 cells. Wild-type mice treated with miR-192 AAV8 had improved glucose sensitivity. Islets isolated from these mice exhibited enhanced GLP-1 potentiated GSIS during perifusion *ex vivo*. These effects were absent in the DIRKO (*Glp1r/Gipr* double knockout) mouse islets, consistent with the idea that miR-192 promotes GLP-1 mediated GSIS through GLP1R. These findings implicate *MIR192* in statin-induced impairment of GSIS by modulating GLP1R, potentially contributing to the susceptibility to NOD in statin users.

## INTRODUCTION

Statins, a class of HMGCR (3-hydroxy-3-methylglutaryl-CoA reductase) inhibitors, are widely prescribed drugs for reducing lipid levels and cardiovascular events and mortality^1^. Despite the well-established benefits, there is extensive evidence that statin use can accelerate the development of new-onset diabetes (NOD)^2^, with up to a 36% increased risk in patients on high-intensity statin therapy^3^. This adverse effect is recognized by the American College of Cardiology, the FDA, and the European Medicines Agency, and has led to changes in statin use guidelines and warning labels on statin prescriptions^4,5^. Recently, genetically proxied HMGCR inhibition was reported to be associated with an elevated risk of type 2 diabetes mellitus (T2DM) in South Asian, East Asian and European populations using Mendelian randomization analyses^6^, consistent with a causal relationship between HMGCR inhibition and diabetes. While it has been emphasized that the new-onset of diabetes does not outweigh the benefits of statins on cardiovascular disease risk^3^, these observations signify the potential for other long-term risks in a relatively high proportion of statin users and highlight the need to elucidate the underlying mechanisms responsible for statin-induced NOD.

A recent study revealed that statins induced hyperglycemia by altering gut microbiota composition and bile acid metabolism, leading to a notable reduction in circulating glucagon-like peptide 1 (GLP-1) levels and disturbed glucose homeostasis^7^. GLP-1, released from enteroendocrine L cells in the intestine in response to a meal, exhibits its incretin activity through the GLP-1 receptor (GLP1R) in β-cells, augmenting glucose-stimulated insulin secretion (GSIS)^8^. In addition to potentiating GSIS, GLP1R signaling also promotes β-cell proliferation and insulin biosynthesis in the pancreas and reduces glucose production, appetite, and gastric emptying in other tissues^9^. Recently developed GLP1R agonists, like semaglutide and liraglutide, have proven to be extremely efficacious drugs for individuals with diabetes, by reducing blood glucose levels and excess body weight and preventing cardiovascular disease, the major cause of mortality in T2DM^10–13^. These benefits point to the potential clinical significance of identifying mechanisms that may augment GLP-1 agonism by upregulating GLP1R.

Multiple studies in β-cell lines and in human and mouse islet cells^14–16^ have demonstrated that statins impair both endogenous and GLP1R mediated insulin secretion. In murine islet cells, this reduction is rapid, reversible, and occurs even with high glucose, and without affecting Ca^2+^ concentrations^14^. Although this effect is well documented, its molecular basis is not clear. While some studies have suggested a role for statin inhibition of isoprenoid synthesis^16^ or mitochondrial function^15^, others have disputed these theories^14^. Thus, additional study is needed to elucidate both the mechanisms by which statin treatment may impair insulin secretion and whether this effect is relevant to developing NOD in statin users.

Here, we employed transcriptomic profiling of patient-derived induced pluripotent stem cells (iPSCs) to gain mechanistic insight into statin-induced NOD. iPSCs retain the genetic characteristics of the donor, exhibit self-renewal, and can be cryopreserved, rendering them a highly versatile cellular model^17,18^. As part of the Pharmacogenomics of Statin Therapy (POST) study, we created a repository of iPSCs derived from statin users who were either susceptible to developing NOD or who maintained stable fasting glucose after statin initiation and discovered a novel molecular factor that regulates GLP1R levels and GLP-1 mediated GSIS and may contribute to statin-induced NOD.

## MATERIALS AND METHODS

### Description of the Kaiser Permanente of Northern California Population (KPNC)

Kaiser Permanente of Northern California (KPNC) is a large group practice-based integrated healthcare in the US. KPNC has well-developed guidelines and standards of care that are uniformly adopted across all sites. In the geographic areas served by Kaiser, approximately 33% of the general population has KPNC coverage. Sociodemographic characteristics are generally representative of the underlying population, except for an under-representation of the extremes of income. For example, the KPNC vs. Bay Area Metropolitan Statistical Area ethnic composition is as follows: white (66% vs. 58%), black (6% vs. 8%), Asian (16% vs. 19%), and other (12% vs. 15%). 19% of the Bay Area Metropolitan Statistical Area self-identifies as Hispanic/Latino, compared to 11% of KPNC members. The 5-year (2009-2013) membership retention rates of persons over ages 40 and 66 are about 65 and 80%, respectively.

### KPNC Electronic Databases

Each Kaiser member is assigned a lifetime unique identifier that is used throughout all databases. Information was drawn from one of three databases:

*<u>KP HealthConnect®</u>* is a fully integrated electronic health record that documents ambulatory visit check-in, hospital-based utilization, clinical documentation of diagnoses, orders for tests and procedures, and prescribed medications. Other functions include documentation of labs, radiology, oncology, and other diagnostic testing, as well as tracking of outside claims and referrals.

KP’s *<u>Pharmacy Information Management System</u>* contains prescription medications dispensed at KPNC hospitals, medical centers, and medical offices for either inpatient or outpatient use. Implemented in all facilities in 1994, the database contains, but is not limited to, medical record number, cost, prescribing practitioner, medicine name, National Drug Code, date of prescription, date medication was dispensed, and dosage and refill information with generation of labels used for dispensing drugs. These data are collected in real time and are considered extremely accurate.

*<u>Laboratory Utilization and Reporting System</u>* captures all ordered and performed laboratory tests from KPNC hospitals, medical centers, and medical offices. The database was created in 1994 and contains, but is not limited to, medical record number, facility code, name of ordering provider, test or procedure name, results, date of test/procedure/result, and abnormal or out-of-range flags.

### Identification of statin users with (NOD susceptible case cohort) or without (NOD resistant control cohort) new-onset diabetes

Using the pharmacy records, we identified individuals who received their first statin prescription between the ages of 40-75 and documented continuous statin use for 3 years. Continuous statin use was defined as having greater than eight 30-day prescription refills per year or at least three 90-day prescription refills per year. Individuals prescribed statin combinations, such as Advicor (niacin/lovastatin) or Vytorin (ezetimibe/simvastatin) as their first prescription were excluded. All statins included in the KPNC formulary are listed in **Table S1**. Individuals were required to have at least two fasting glucose measures in the 3 years prior to the start of statin treatment. The range of these values for each individual had to be within 30 mg/dL, and all values had to fall between 50-110 mg/dL (**Fig. S1**).

The following criteria within 3 months to 3 years after statin initiation were used to identify and recruit NOD susceptible cases: 1) two or more fasting glucose ≥ 126 mg/dL, 2) one fasting glucose ≥ 126 mg/dL plus a diabetes diagnosis based on ICD (International Classification of Diseases) codes (**Table S2**), 3) one fasting glucose ≥ 126 mg/dL plus any prescription of glucose lowering drugs, or 4) a fasting glucose increase of ≥ 13 mg/dL (**Fig. S1A**). Individuals with diabetes ICD codes prior to their first statin prescription or within the first 3 months of statin use were excluded. Additionally, anyone with a fasting glucose measurement ≥ 126 mg/dL within the first 3 months on statin was excluded. Any prescription for glucose lowering drugs such as anti-hyperglycemic agents (alpha-glucosidase inhibitors, dipeptidyl peptidase-4 inhibitors, meglitinides, sulfonylureas, biguanides, and thiazolidinediones) and insulin (**Table S3**) in the 3 years before and 3 months after statin initiation was also an exclusion criterion. Furthermore, individuals were excluded who had prescriptions for drugs that may raise glucose, such as oral corticosteroids (**Table S4**) or who had ICD/CPT codes for bariatric surgery (**Table S5**), in the 3 years before or 3 years after starting statin treatment.

NOD resistant controls were statin users with at least three fasting glucose measures during the first 5 years of treatment, all of which had to be between 50-110 mg/dL (**Fig. S1B**). Individuals with a diabetes or bariatric surgery ICD code or use of glucose-modifying drugs (**Tables S2-S5**) during the 3 years prior to and the first 5 years of statin treatment were excluded from the NOD resistant control cohort.

Individuals who met the specified criteria were contacted and recruited for the Pharmacogenomics of Statin Therapy (POST) study. A total of 185 statin-induced NOD susceptible cases and 320 NOD resistant controls consented to participate. Written informed consent was obtained from all participants, and the study was conducted with IRB approval, including authorization to access protected health information within the electronic health record.

### Creation and authentication of iPSCs

iPSCs were reprogrammed from CD34^+^ peripheral blood mononuclear cells (PBMCs) isolated from blood samples of study participants and validated as previously described^19^. Briefly, CD34^+^ PBMCs were expanded, nucleofected with episomal vectors of POU5F1 (aka OCT-4), SOX2, KLF4, L-MYC, LIN28, EBNA1, and shRNA for TP53^20^, and seeded onto mitomycin C treated SNL feeder cells. After emergence of iPSC colonies, TRA-1-60 expressing cells were selected using Magnetic Activated Cell Sorting to obtain one pooled cultured iPSC line per study participant. Expression levels of pluripotency markers POU5F1 or TRA-1-60, and differentiation marker SSEA-1 in iPSCs were visualized with immunohistochemistry and quantified with flow cytometry. Representative lines were sent for KaryoStat, PluriTest (Thermo Fisher Scientific), or karyotype (Cedar Sinai RMI iPSC Core) analyses. The pluripotency of certain iPSCs was further confirmed by positive differentiation of representative lines into endoderm, mesoderm and ectoderm using the STEMdiff Trilineage Differentiation Kit (StemCell Technologies, #05230) followed by immunostaining and flow cytometry analyses using a BD LSR Fortessa flow cytometer at the UCSF MLK Cores Research Facility^19^.

### Cell culture

The INS-1 rat insulinoma cell line was maintained in RPMI 1640 media with 10% fetal bovine serum (FBS), 1 mM sodium pyruvate, 10 mM HEPES, and 55 μM 2-Mercaptoethanol. The βTC3 mouse insulinoma cell line was maintained in DMEM containing 25 mM glucose, 1mM pyruvate, 2mM GlutaMax, 15% heat-inactivated horse serum, and 2.5% FBS. HEK293T cells were maintained in DMEM containing 25 mM glucose, 10% FBS, and 1% penicillin/streptomycin. All cell lines were kept at 37 °C in a humidified incubator containing 5% CO_2_.

iPSCs were cultured on plates coated with Cultrex reduced growth factor basement membrane (Trevigen # 3533-001-02), fed daily with mTeSR1 (StemCell Technologies # 85850), passaged routinely using ReLeSR (StemCell Technologies # 05872), and kept at 37 °C in a humidified hypoxia incubator with 5% O_2_ and 5% CO_2_.

Primary mouse islets were isolated as previously described^21^. Briefly, after mice were sacrificed with isofluorane and cervical dislocation, the common bile duct was clamped off at the duodenum and collagenase (7.5 mg/mL, Sigma, #C7657) in HBSS (5 mM glucose, 1 mM MgCl_2_) was injected into the common bile duct. The pancreas was removed and digested at 37 °C for 12 minutes. Digested tissue was washed with ice-cold HBSS (5 mM glucose, 1mM MgCl_2_, 1 mM CaCl_2_) 3 times. Islets were separated by Histopaque (Sigma, #10771) gradient and hand-picked in complete RPMI (10% FBS, 1% penicillin/streptomycin).

### RNA-seq

iPSCs from 24 NOD susceptible cases and 24 NOD resistant controls were exposed to 250 nM atorvastatin, 500 nM simvastatin, or a mock buffer for 24 hours before total RNA was isolated using RNeasy Mini Kit (Qiagen, #74106). PolyA-selected, strand-specific paired-end RNA-seq libraries were prepared at the Northwest Genomics Center and sequenced using Illumina technology. Sequences were aligned to the human (GRCh38) genome and transcriptome (GENCODE) using STAR^22^, fragment counts aligning to genes were quantified using FeatureCounts^23^, and gene expression levels were adjusted for library size and variance stabilized (∼log_2_ transformation) using DESeq2^24^. Statin-induced changes in gene expression were calculated as statin-mock variance stabilized deltas for each statin type, and differential changes in gene expression between NOD susceptible cases and NOD resistant controls were identified using t-tests. The RNA-seq data is being deposited into dbGaP under the POST study ID phs003596.

### Transfection of miRNA mimic and inhibitor

INS-1 and βTC3 cells were transfected with mirVana miRNA mimic (Invitrogen, #4464066) for hsa-miR-192-5p (Assay ID MC10456) or hsa-miR-204-5p (Assay ID MC11116), mirVana miRNA inhibitor (Invitrogen, #4464084) for hsa-miR-192-5p (Assay ID MH10456) or hsa-miR-204-5p (Assay ID MH11116), or equal concentration of negative control mirVana miRNA mimic (Invitrogen, #4464058) negative control inhibitor (Invitrogen, #4464076) using Lipofectamine RNAiMAX (Invitrogen, #13778075).

Primary islets were dispersed by trypsin (0.25% in Trypsin/EDTA) digestion for 15 minutes at 37 °C and transfected with either mirVana miRNA mimic (Invitrogen, #4464066, Assay ID MC10456) or negative control mirVana miRNA mimic (Invitrogen, #4464058) using Lipofectamine RNAiMAX (Invitrogen, #13778075) in Optimem media and shaken gently overnight in the incubator (37 °C, 5% CO_2_). Twelve to eighteen hours after transfection, 2X RPMI media (20% FBS, 2% penicillin/streptomycin) was added to the plates. Forty-eight hours after transfection, islet cells from 4-5 mice were pooled and reaggregated into pseudoislets per manufacturer’s instructions (AggreWell™400, StemCell Technologies). After harvesting, pseudoislets were incubated in complete RPMI overnight before perifusion.

### RNA isolation and qPCR

Total RNA was isolated from cells with mirVana miRNA Isolation Kit (Thermo Fisher Scientific, #AM1560) or Quick-RNA Miniprep Kit (Zymo, #R1054) 48 hours after transfection. The cDNA was prepared from 10 ng of total RNA from cells with a TaqMan advanced miRNA cDNA synthesis kit (Thermo Fisher Scientific, #A28007). Real-time PCR with TaqMan advanced miRNA assays (Thermo Fisher Scientific, #A25576) was used to quantify mature miRNA in the samples: hsa-miR-192-5p (478262_mir), hsa-miR-204-5p (478491_mir), or hsa-miR-16-5p (477860_mir). Reverse transcription was done with High-Capacity cDNA Reverse Transcription Kit (Fisher Scientific, #43-688-14). *Glp1r* (Rn00562406_m1 or Mm00445292_m1) and *Clptm1* (Rn01457493_m1 or Mm01321458_m1) transcript levels were determined by qPCR on a QuanStudio 6 Pro (Thermo Fisher Scientific).

For primary mouse islets, miRNAs were isolated per TaqMan® Advanced miRNA Assay (Thermo Fisher Scientific, #A25576) instructions and reverse transcription per TaqMan® Advanced miRNA cDNA Synthesis Kit (Thermo Fisher Scientific, #A28007). RT-qPCR was performed on QuantStudio 5 and miR-192 (Thermo Fisher 478262_mir, hsa-miR-192-5p) and miR-16 (Thermo Fisher 477860_mir, hsa-miR-16-5p) expression were quantified as fold change compared to controls. mRNA was isolated with TRIzol reagent per manufacturer’s instructions. Reverse transcription was done with High-Capacity cDNA Reverse Transcription Kit (Fisher Scientific, #43-688-14). mRNA abundance of *Glp1r* (Thermo Fisher, Mm00445292_m1) and *Actb* (Thermo Fisher, Mm00607939_s1) were determined with QuantStudio 5 and quantified as fold change compared to controls.

### Western blots

Protein extracts were prepared using RIPA buffer (Thermo Fisher Scientific, #89900) with Halt protease inhibitor cocktail (Thermo Fisher Scientific, #78429), and protein concentration was determined with Pierce BCA protein assay (Thermo Fisher Scientific, #23227). Equal amounts of protein samples were separated by electrophoresis on a 4% to 20% Tris-Glycine gel (Thermo Fisher Scientific, #XP04202BOX) and transferred to a PVDF membrane. Antibodies used in the current study include anti-GLP1R (Abcam, #ab218532, 1:1250), anti-GAPDH (Santa Cruz Biotechnology, #sc-166545, 1:1400), goat-anti-rabbit IgG (Thermo Fisher Scientific, #31460, 1:6000), and m-IgGκ BP-HRP (Santa Cruz Biotechnology, #sc-516102, 1:6000).

### 3’ UTR reporter constructs

The LightSwitch luciferase reporter construct, pLS-GLP1R-3’UTR, was purchased from SwitchGear Genomics (#S810047), and encompasses bases 1411-3187 of the GLP1R transcript NM_002062.5. The following pairs of primers were used to mutate the predicted miR-192 binding site AGGTCAA at NM_002062.5 position 1813-1819 to GTAGTGC: 5’-GTGCCGGCTTATTAGTGAAACTGGGGCTTG-3’ and 5’-TACTGAGTTTGAGTCTGGGGTTGATTTGCGGC-3’. An empty LightSwitch 3’UTR reporter vector (SwitchGear Genomics, #S890005) was used as a control construct in luciferase reporter assays. The positive control SCD 3’ UTR reporter construct was a kind gift from Dr. Vesa M. Olkkonen.

### Luciferase reporter assay

Cells were plated in white 96-well plates overnight and co-transfected for 24-48 hours with each luciferase reporter construct and miR-192-5p mimic, inhibitor, or non-targeting negative controls using DharmaFECT DUO transfection reagent (Dharmacon, #T-2010-02) following manufacture’s instruction. Luciferase activity was measured with LightSwitch assay reagents (SwitchGear Genomics, #LS100) on a Synergy H1 microplate reader.

### Insulin secretion assays

INS-1 cells were plated in 96-well plates and transfected the next day with various miRNAs for 48 hours. For the statin study, transfected cells were treated 24 hours after transfection with either 500 nM or 1000 nM of simvastatin or a mock buffer for 24 hours. On the day of the insulin secretion assay, transfected INS-1 cells were pre-incubated in KREBS-Ringer Bicarbonate (KRB) buffer containing 1% BSA for 30 min at 37 °C and then incubated with 3 or 17.8 mM glucose and 50 nM GLP-1 in KRB buffer containing 1% BSA for 30 min. The insulin containing buffer was then collected from each well and centrifuged at 450 x g for 5 min. The resulting supernatant was stored at −20 °C until ready for insulin measurement, whereas the cells were washed with DPBS and lysed in CelLytic M for protein quantification using a Bradford protein assay. Insulin concentrations were measured using a rat/mouse insulin ELISA kit (Millipore, #EZRMI-13K) following the manufacturer’s instructions.

Perifusion studies were performed by placing 80-100 primary mouse islets (or pseudoislets from transfected islets after aggregation) in a Biorep (Biorep Technologies) perifusion chamber and stimulated with low glucose (2.8 mM, equilibration 40 minutes), very high glucose (16.7 mM), and high glucose (10 mM) with and without GLP-1 (0.3 μM, Bachem #4034865), GIP (100 nM, D-Ala2-GIP, Bachem #4054476), and KCl (30 mM) in KRBH (115 mM NaCl, 5 mM KCl, 2.5 mM CaCl2, 24 mM NaHCO3, 10 mM HEPES, and 1% BSA; pH = 7.4). The flow rate was 100 μl/minute, and the chamber temperature was 37 °C. Perifusion effluents were stored at -80°C until insulin quantification with ALPCO High Range Insulin ELISA (80-INSMSH).

### Animal studies

Animal studies were approved by either the Institutional Animal Care and Use Committee at the University of California San Francisco or the Animal Care Committee (UOHI-AUP-2909) in accordance with the Canadian Guide for Care and Use of Laboratory Animals. Mice were group housed in standard cages in climate-controlled facilities on a 12-hour light/dark cycle and given ad libitum access to food and water unless otherwise indicated.

Wild-type C57BL/6J male mice were put on a GAN diet (Research Diets, #D09100310, 40 kcal% fat, 20 kcal% fructose and 2% cholesterol) at 7-week-old for 4 weeks before receiving either miR-192 (AAV8-EF1α-mmu-mir-192-eGFP, Vector Biolabs) or control (scAAV8-EF1α-ctrl-miR-eGFP, Vector Biolabs) AAV vectors (8×10^12^ viral genomes/kg) via intraperitoneal injection. Four more weeks later, an intraperitoneal glucose tolerance test was performed with 1.5 g/kg glucose after a 6-hour fast, and plasma glucose levels were determined over a 2-hour period. Additional cohorts of male C57BL/6J wild-type or double incretin receptor knockout (DIRKO)^25^ mice were fed a high fat diet (HFD; TD88137, 42 kcal% fat and 0.2% cholesterol) starting at 12 weeks of age. After 8 weeks on HFD, mice received a tail vein injection of a miR-192 (AAV8-EF1α-mmu-mir-192-eGFP, Vector Biolabs) or control (scAAV8-EF1α-ctrl-miR-eGFP, Vector Biolabs) AAV (1 × 10^11^ GC/mouse in 200 µl). To quantify glucose tolerance, mice were fasted for 5 hours, injected with glucose solution (2 g/kg) intraperitoneally and blood glucose was measured by glucometer both prior to glucose injection and every 15 minutes for 90 minutes post-injection.

## RESULTS

### Generation of iPSCs from NOD susceptible cases versus NOD resistant controls

Peripheral blood mononuclear cells (PBMCs) were collected from cohorts of NOD susceptible cases and NOD resistant controls in the Pharmacogenomics of Statin Therapy (POST) study (**Fig. S1**). PBMC-derived iPSC lines were generated from 24 NOD susceptible cases and 24 NOD resistant controls using methods we previously described^19^. Immunohistochemistry and flow cytometry were used to verify iPSC marker expression, with greater than 90% of cells positive for the pluripotency markers TRA-1-60 or POU5F1 (aka OCT-4), and fewer than 5% positive for the differentiation marker SSEA-1 (data not shown).

The demographic and clinical characteristics of the 48 donors are summarized in **Tables 1, S6-S7**. Prior to statin initiation, both groups had similar plasma total and LDL cholesterol levels. However, pre-statin triglyceride levels trended higher in the NOD susceptible cases (174.96 ± 128.61 vs 123.79 ± 58.70 mg/dL, *p* = 0.086). Average baseline fasting glucose was significantly higher in the NOD susceptible cases (99.54 ± 6.45 vs 94.00 ± 5.94 mg/dL, *p* = 0.004). The magnitudes of statin-induced changes in total cholesterol, LDL cholesterol, and triglyceride levels did not differ significantly between the two groups. As expected, the NOD susceptible cases exhibited significantly greater statin-induced fasting glucose changes compared to the NOD resistant controls.

**Table 1.**
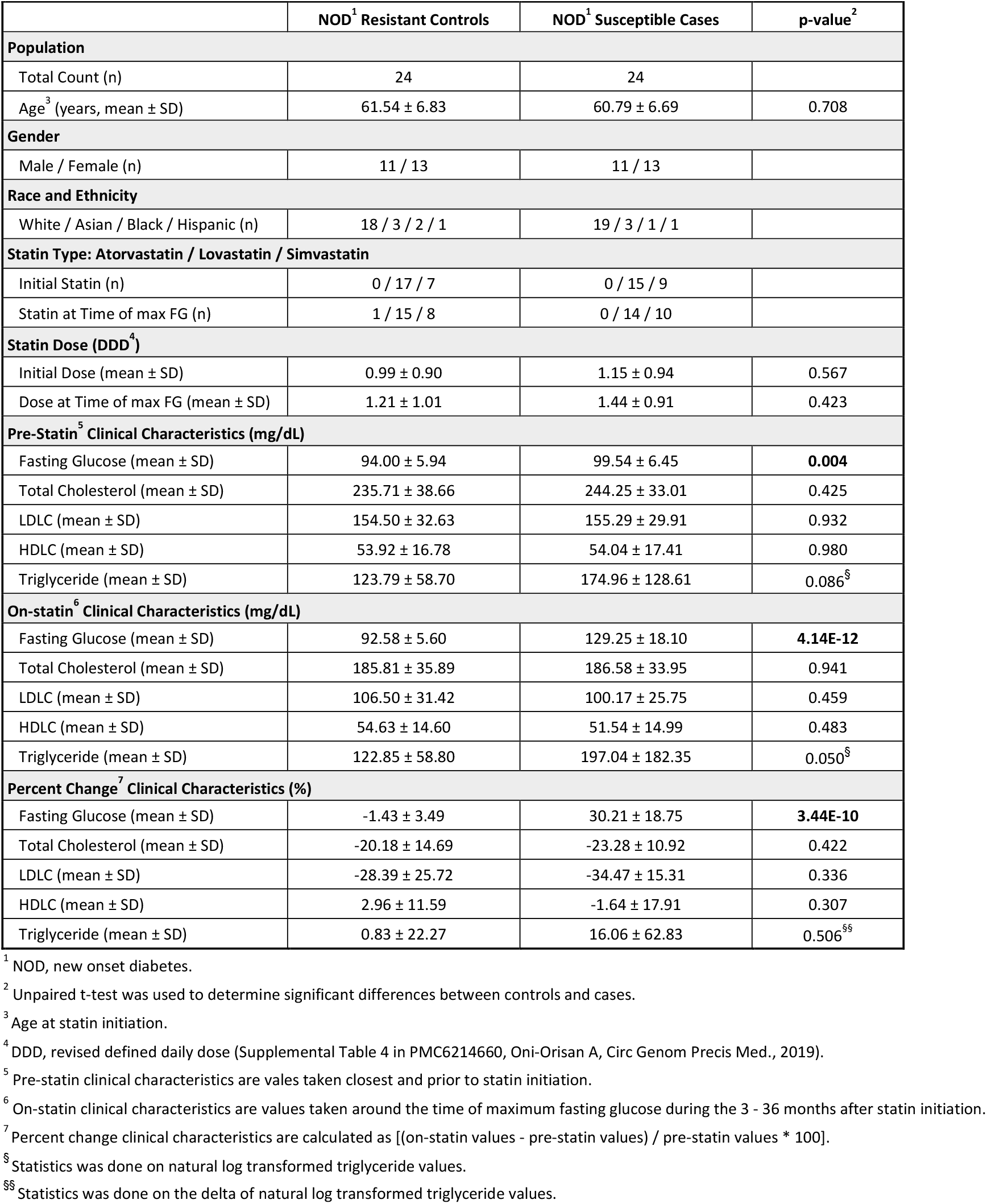
Demographic and Clinical Characteristics.

### Identification of MIR194-2HG as a putative factor underlying statin-induced dysglycemia

We performed RNA-seq on iPSCs from NOD susceptible cases and NOD resistant controls (n=24/group) incubated with 250 nM atorvastatin, 500 nM simvastatin, or a mock buffer for 24 hours. These statin concentrations were determined using dose-response analyses as the lowest concentrations that generated a robust and reproducible increase in *HMGCR* and decrease in *MYLIP*, well-known effects of statin^26,27^ (**Fig. S2**). The different doses of atorvastatin and simvastatin accounts for their relative LDL cholesterol lowering potency^28^. From RNA-seq analysis, the *MIR194-2 host gene* (*MIR194-2HG*) on chromosome 11 emerged as a transcript of interest, as it was among the most differentially changing transcripts between the two groups that also exhibited opposite directionality of statin-induced changes between case and control lines (**Table S8**). Specifically, exposure to either simvastatin or atorvastatin increased *MIR194-2HG* expression in the NOD resistant controls, but reduced *MIR194-2HG* in the NOD susceptible cases (*p* < 0.01, unpaired t-test, **Fig. 1A**). In contrast, statin treatment increased *HMGCR* (**Fig. 1B**) and decreased *MYLIP* (**Fig. 1C**) gene expression as expected in both groups. In a subset analysis of iPSCs from donors who met the strict normal fasting glucose levels (< 100 mg/dL) prior to statin initiation (19 NOD resistant controls and 13 NOD susceptible cases), we observed similar trends in the differential regulation of *MIR194-2HG* by statins (**Fig. S3**).

**Figure 1.**
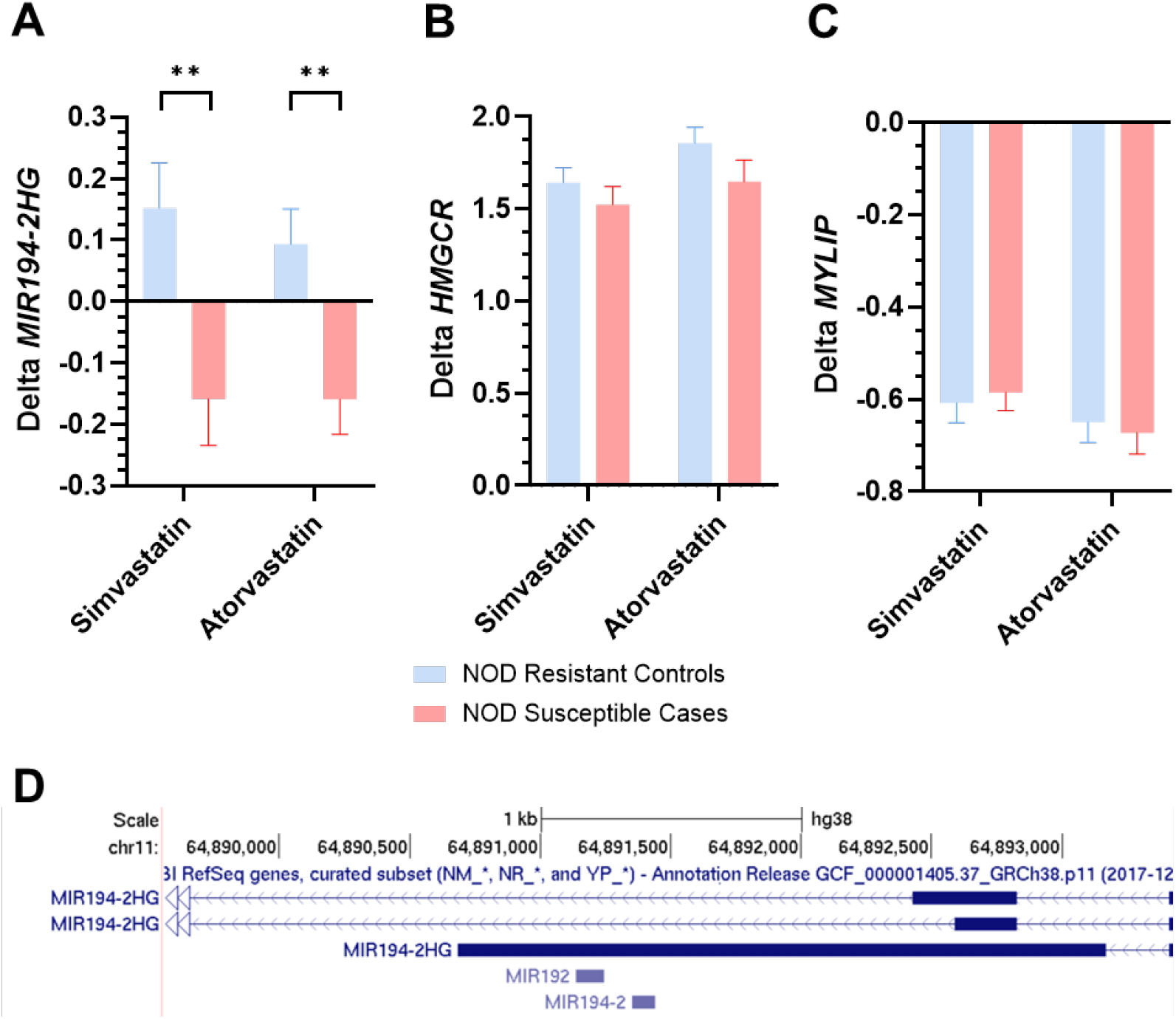
Identification of *MIR194-2HG* as a putative factor underlying statin-induced new-onset Type 2 diabetes. **(A-C)** Change in transcript levels in iPSCs derived from statin users categorized as NOD susceptible cases (n=24) or NOD resistant controls (n=24) after 24hr incubation with 500 nM simvastatin, 250 nM atorvastatin or mock control buffer. Values were variance stabilized (∼log2 transformation) in DESeq2 and deltas were calculated as statin-control expression levels. Data are presented as mean +/- SEM, significance was determined by unpaired t-test, \*\**p* < 0.01. **(D)** Schematic of the *MIR194-2HG*, which encompasses two miRNA genes, *MIR192* and *MIR194-2*.

*MIR194-2HG* encodes two miRNA genes, *MIR192* and *MIR194-2* (**Fig. 1D**). Notably, *MIR194-2HG* is the only source of *MIR192*, whereas *MIR194* can also be transcribed from chromosome 1 (*MIR194-1)*. Elevated levels of circulating miR-192-5p, the mature microRNA derived from the 5’ arm of *MIR192*, have been observed in individuals with diabetes (type 1 or 2) or impaired fasting glucose compared to those with normal fasting glucose^29–32^. Therefore, we focused additional studies on miR-192-5p, which we will refer to as miR-192 hereafter.

### miR-192 increases GLP1R levels in β-cell models

miR-192 has been shown to downregulate GLP1R in the human renal tubular epithelial cell line HK-2^33^ and the human enteroendocrine cell line NCI-H716^34^. miR-192 exhibits tissue-restricted expression but is present in the human pancreas, islets and β-cells^35,36^. To test whether miR-192 regulates GLP1R in pancreatic β-cells, we transfected INS-1 cells, a rat pancreatic β-cell line, with a miR-192 mimic or a non-targeting control. Surprisingly, *Glp1r* transcript levels were upregulated 1.41 ± 0.03-fold by the miR-192 mimic (**Fig. 2A-B**), an effect that was confirmed in the mouse β-cell line βTC3 (**Fig. S4**). This induction was reversed upon addition of a miR-192 inhibitor (**Fig. 2A-B**), while treatment with the miR-192 inhibitor alone reduced *Glp1r* transcript levels (0.84 ± 0.01-fold, **Fig. 2C-D**). Since miRNAs typically downregulate mRNAs, the unexpected upregulation of *Glp1r* by miR-192 prompted us to treat INS-1 cells with a miR-204-5p mimic, previously reported to target *Glp1r*^37^, and observed the expected reduction in *Glp1r* levels (**Fig. 2E**). The miR-192 mimic did not increase transcript levels of glucose-dependent insulinotropic polypeptide (GIP) receptor (*Gipr*), another G-protein coupled receptor involved in insulin secretion (**Fig. S5**).

**Figure 2.**
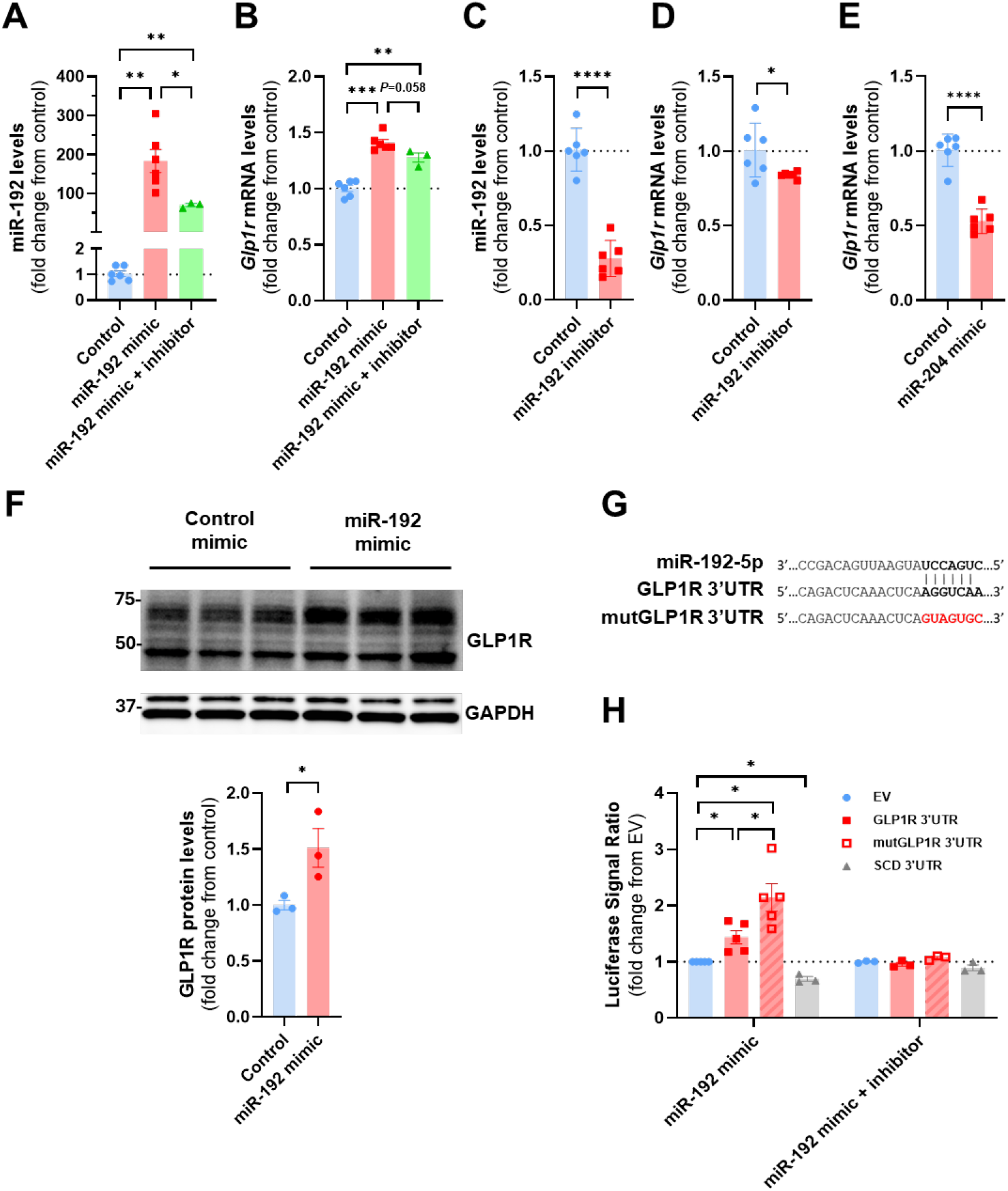
miR-192 mimic upregulates GLP1R in INS-1 cells and miR-192 inhibitor reversed the effect. **(A**,**B)** INS-1 cells were transfected with 10 nM miR-192 mimic, 10 nM miR-192 mimic plus 10 nM miR-192 inhibitor, or non-targeting controls for 48 hours, and transcript levels of miR-192 **(A)** and *Glp1r* **(B)** were quantified. **(C**,**D)** INS-1 cells were transfected with 30 nM miR-192 inhibitor or a non-targeting control for 48 hours, and transcript levels of miR-192 **(C)** and *Glp1r* **(D)** were quantified. **(E)** INS-1 cells were transfected for 48 hours with 10 nM miR-204-5p mimic or a non-targeting control, and *Glp1r* transcript levels were quantified. For panels A-E, *Glp1r* transcript levels were normalized to *Clptm1*, miR-192 levels were normalized to miR-16-5p, and all transcripts were quantified by qPCR **(F)** INS-1 cells were transfected with 10 nM miR-192 mimic or a non-targeting control for 48 hours, and GLP1R protein levels normalized to GAPDH were quantified by western blot analysis. **(G)** A schematic of the sequence predicted to be targeted by miR-192 in unedited and mutant human *GLP1R* 3’UTR luciferase constructs. **(H)** INS-1 cells were co-transfected with luciferase constructs containing part of the human *GLP1R* 3’ UTR either unedited or after mutagenesis (mutGLP1R3’UTR) of the predicted miR-192 binding site, SCD 3’ UTR, or an empty vector (EV), along with 10 nM miR-192 mimic, 10 nM miR192 mimic plus 10 nM miR-192 inhibitor, or non-targeting controls. Luciferase signal ratios for miR-192 mimic over non-targeting control in cells transfected with different plasmids were compared to those in cells transfected with EV. Data are presented as mean +/- SEM, statistical significance was determined by unpaired t-test, \**p* < 0.05, \*\**p* < 0.01, \*\*\**p* < 0.001, \*\*\*\**p* < 0.0001.

We next confirmed that the miR-192 mimic increased GLP1R protein levels by 1.51 ± 0.17-fold using western blot analysis (**Fig. 2F, S6A-B**). Since many GLP1R antibodies have poor specificity^38^, we used a commercially available antibody (Abcam, # ab218532) thoroughly validated in mouse pancreatic islets, cellular models, and rat tissues using immunohistochemistry staining and western blot^39,40^. To further confirm the antibody specificity, we assayed HEK293T cells and found no detectable GLP1R protein (as expected) and reduced GLP1R protein abundance in INS-1 cells treated with a *Glp1r* siRNA compared to scrambled control (**Fig. S6C-E**).

To examine whether miR-192 upregulates GLP1R through the 3’ UTR, we cotransfected INS-1 cells with a luciferase reporter containing 1777 bp of the human *GLP1R* 3’ UTR with the miR-192 mimic or a non-targeting control. Prior reports identified a miR-192 binding site in the *GLP1R* 3’ UTR at position 1813-1819 of the *GLP1R* transcript NM_002062.5 (**Fig. 2G**)^33,34^. The miR-192 mimic significantly increased the luciferase signal by 1.44 ± 0.12-fold in cells transfected with the *GLP1R* 3’ UTR compared to the empty 3’ UTR vector (**Fig. 2H**). Notably, miR-192 increased the luciferase signal even after disruption of the reported binding site (mutGLP1R 3’UTR, **Fig. 2G-H**). As a positive control, we tested the effect of the miR-192 mimic on a luciferase construct containing the 3’ UTR of *SCD*, a direct target of miR-192^41^, and found the expected reduction in luciferase levels (**Fig. 2H**). miR-192 effects on luciferase levels with either the *GLP1R* or *SCD* 3’ UTR vectors were reversed upon the addition of the miR-192 inhibitor (**Fig. 2H**).

### miR-192 increases GLP-1 augmented GSIS and rescues statin-induced impairment of GSIS in INS-1

Based on the increase in GLP1R levels, we hypothesized that miR-192 may enhance the effects of GLP-1 on GSIS. Media insulin levels were elevated after stimulation with 17.8 mM glucose plus 50 nM GLP-1 or 25 nM exendin-4 (Ex4, a GLP1R agonist) in INS-1 cells transfected with miR-192 mimic compared to control (**Fig. 3A**). In contrast, there was no effect of miR-192 on insulin levels with non-stimulatory glucose concentrations (3 mM), or after the addition of a GLP1R antagonist (exendin 9-39, or Ex9) (**Fig. 3A**). Prior reports have shown that statin treatment can inhibit insulin secretion^14–16^. Indeed, simvastatin treatment dose-dependently reduced GLP-1 mediated GSIS in INS-1 cells transfected with non-targeting controls (**Fig. 3B**). Notably, the reduction in media insulin observed in simvastatin-treated INS-1 cells was rescued by miR-192 overexpression, while the addition of the miR-192 inhibitor reversed this effect (**Fig. 3B**).

**Figure 3.**
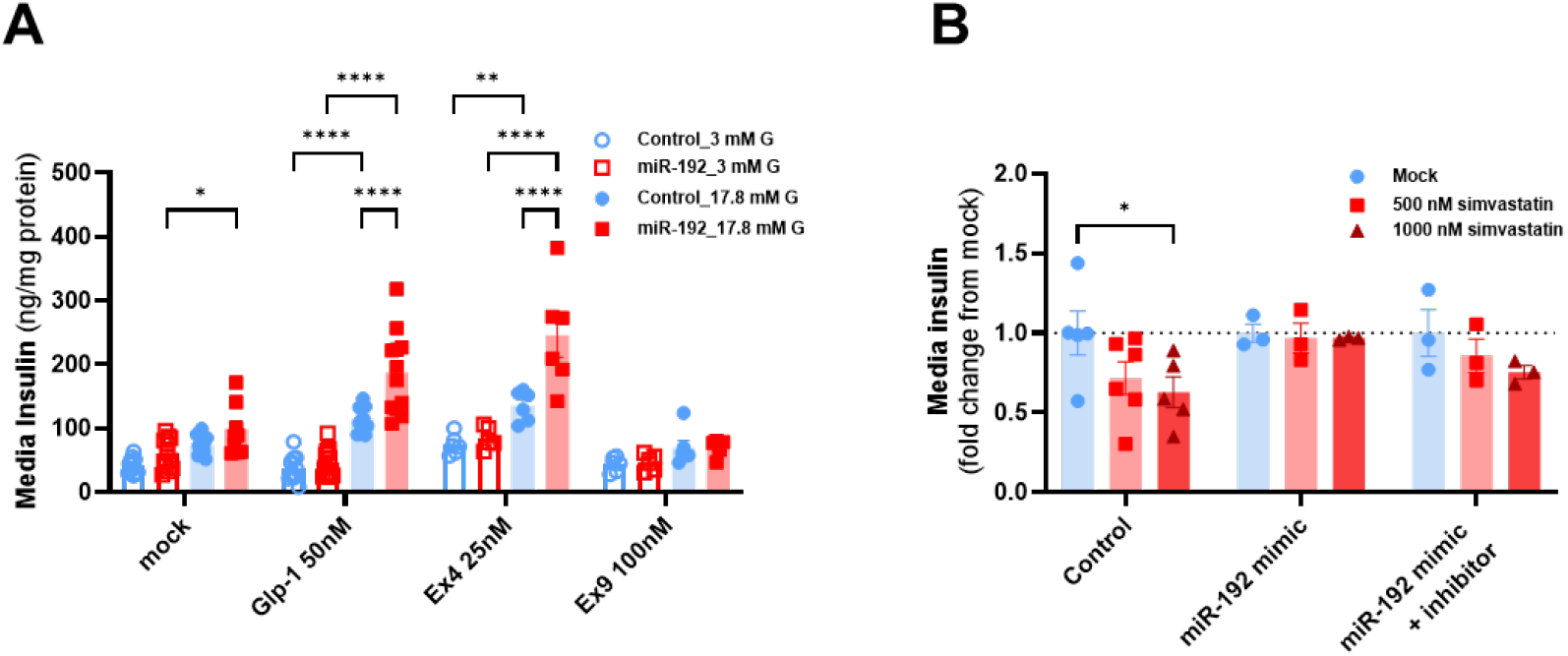
miR-192 mimic increases Glp-1 augmented glucose-stimulated insulin secretion (GSIS) in INS-1 and protects against statin-induced reductions in GSIS. **(A)** INS-1 cells were transfected with the miR-192 mimic or scrambled control for 48 hours, stimulated for 30 minutes with glucose (3 mM or 17.8 mM) and a GLP1R agonist (50 nM GLP-1 or 25 nM exendin-4) or GLP1R antagonist (100 nM exendin 9-39), and insulin in the media was quantified by ELISA, n=6-12/treatment. **(B)** INS-1 cells were transfected with the miR-192 mimic, mimic + inhibitor, or scrambled controls for 24 hours, after which cells were treated with simvastatin or a mock buffer for 24 hours. GSIS was then performed with 17.8 mM glucose + 50nM GLP-1 in corresponding simvastatin or mock buffer. After 30 minutes media insulin was quantified. Values were calculated as a fold change from the mock treatment. Data are presented as mean +/- SEM, and significance was determined by two-way ANOVA adjusted for multiple comparisons, \**p* < 0.05, \*\**p* < 0.01, \*\*\*\**p* < 0.0001.

### miR-192 upregulates GLP-1 augmented islet GSIS ex vivo and may improve glucose tolerance in vivo

To evaluate the effects of miR-192 *in vivo*, wild-type C57BL/6J male mice were fed a GAN diet for 4 weeks after which animals were injected once intraperitoneally with a miR-192 versus a scrambled control AAV8. After an additional 4 weeks, the miR-192 AAV8 treated animals had significantly improved glycemic control during an intraperitoneal glucose tolerance test (IpGTT) compared to control-treated mice (**Fig. S7**). We next tested the effects of miR-192 in HFD-fed male wild-type C57BL/6J mice and DIRKO (*Glp1r/Gipr* double knockout^25^) mice. Four weeks after a one-time tail vein injection of miR-192 or control AAV8 vectors, miR-192 treated wild-type mice trended toward lower weight gain (unpaired t-test *p* = 0.063, **Fig. S8A**), improved glucose tolerance via IpGTT (*p* = 0.0675, **Fig. 4A-B**), and displayed higher total insulin secretion during a perifusion assay in isolated islets (**Fig. 4C-D**). These effects were not seen in the DIRKO mice (**Fig. S8A, 4C-D**). Closer evaluation of the dynamic insulin secretion assay revealed that compared to islets from control-treated animals, the islets from the miR-192 treated wild-type animals had higher insulin secretion in response to 10 mM glucose plus 0.3 μM GLP-1 or 100 nM GIP (**Fig. 4F-G**), but not to 16.7 mM glucose (**Fig. 4E**) or KCl (**Fig. 4H**). In wild-type animals, miR-192 and *Glp1r* transcripts were insignificantly trending to higher levels in miR-192 islet samples (**Fig. S8B-C**). *Glp1r* mRNA was undetectable in DIRKO islet samples (**Fig. S8C**). Lastly, in the miR-192 treated wild-type animals, there was a statistically significant correlation between islet *Glp1r* mRNA levels and total perifusion AUC (**Fig. 4I**).

**Figure 4.**
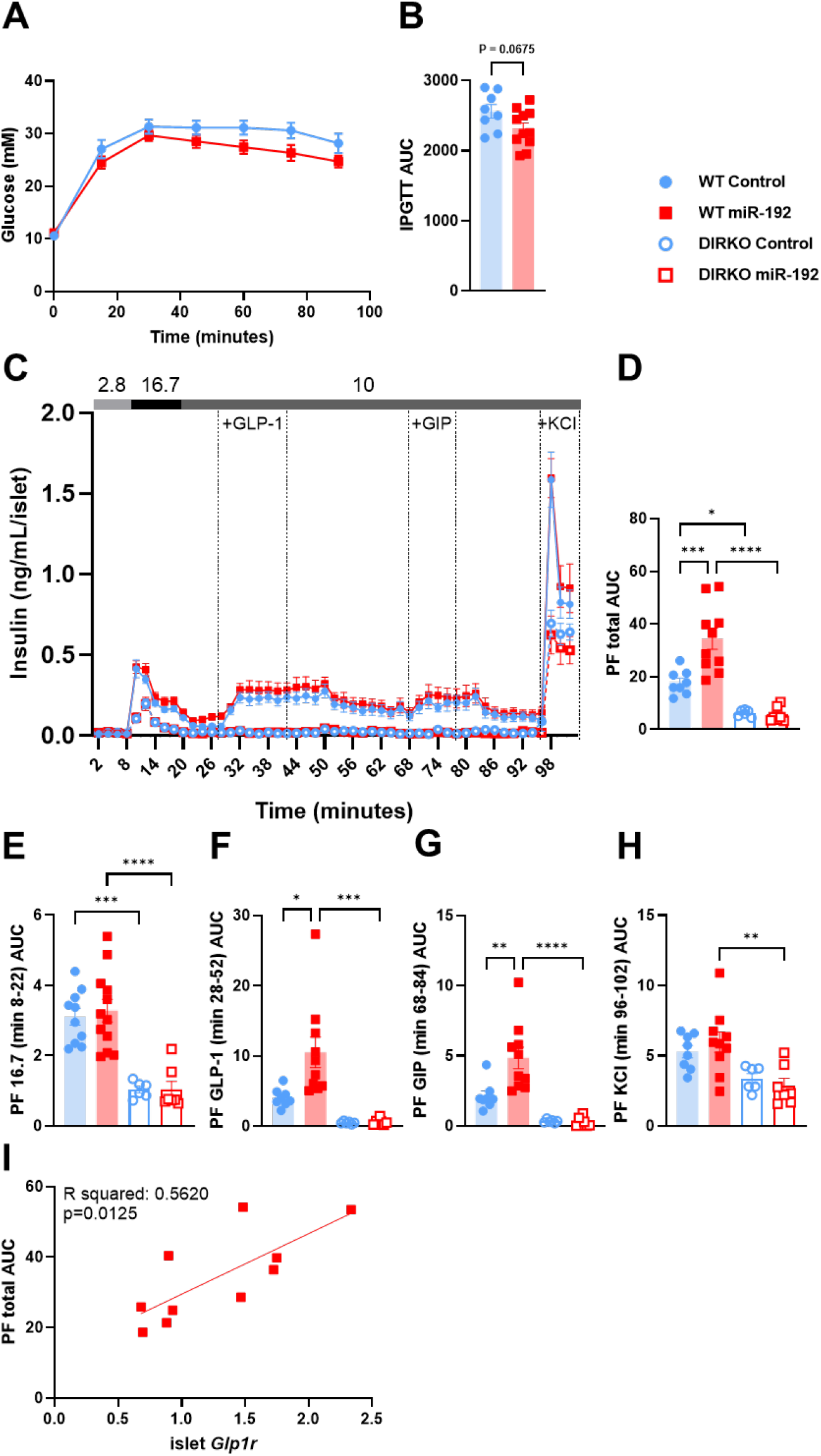
miR-192 overexpression *in vivo*. t 12 weeks of age, male C57BL/6J wild-type and double incretin receptor knockout (DIRKO) mice were placed on a high-fat diet (42 kcal% fat, 0.2% cholesterol). Following 8 weeks of dietary exposure, mice were administered once via tail vein injection with either miR-192 or a scrambled control AAV8 construct. Intraperitoneal glucose tolerance test was performed 4 weeks after the AAV treatment. **(A)** Blood glucose levels during i.p. glucose tolerance test (2 g/kg) in wildtype mice. **(B)** Area under the curve (AUC) glucose during i.p. glucose tolerance test (2 g/kg) in wildtype mice. **(C)** Insulin measured during dynamic perifusion (PF) experiment of wildtype and DIRKO mouse islets. **(D)** Total AUC insulin during perifusion experiment shown in (C). **(E-H)** Perifusion incremental AUC during treatment of 16.7 mM glucose **(E)**, 10 mM glucose and 0.3 μM GLP-1 **(F)**, 10 mM glucose and 100 nM GIP **(G)**, or 10 mM glucose and 30 mM KCl **(H)** in wildtype and DIRKO mouse islets. **(I)** Correlation analysis of islet *Glp1r* transcript levels normalized to *ActB* and total perifusion AUC in wildtype miR-192 mice. Data are presented as mean +/- SEM, significance was determined by unpaired t-test (B), one way ANOVA adjusted for multiple comparisons (D-H), or correlation analysis (I). \**p* < 0.05, \*\**p* < 0.01, \*\*\**p* < 0.001, \*\*\*\**p* < 0.0001. Abbreviation: PF, perifusion.

Finally, as an alternative model, we tested the effects of miR-192 overexpression in primary murine islets transfected *ex vivo*. Islets from wild-type or DIRKO mice were dispersed into single cells, transfected with either miR-192 mimic or a non-targeting control, and reaggregated into pseudoislets. The miR-192 mimic increased transcript levels of both miR-192 and *Glp1r* transcripts in wild-type islets (**Fig. S9A**), with a very similar magnitude of effect on *Glp1r* as seen in the INS-1 cells (**Fig. 2B**). miR-192 overexpression was also achieved in DIRKO islets with no detectable *Glp1r* transcripts as expected (**Fig. S9B**). We confirmed that pseudoislets displayed distinct first and second phases of insulin secretion with high glucose (16.7mM), and a transient and discrete increase upon the addition of 0.3 μM GLP-1 (**Fig. S9A**). Notably, we observed a sustained increase in GLP-1 augmented GSIS in the miR-192 treated islets from wild-type animals (**Fig. S9A**), while there was no effect of miR-192 in DIRKO pseudoislets from the DIRKO mice (**Fig. S9B**). These findings agree with the AAV8 animal studies and together suggest that miR-192 elevates GLP-1 potentiated GSIS through up-regulation of GLP1R and may improve glucose tolerance *in vivo*.

## 4. DISCUSSION

Using our unique resource of iPSCs from statin users who had worsened fasting glucose measures on statin (NOD susceptible cases) versus those whose fasting glucose remained stable despite statin treatment (NOD resistant controls), we have identified *MIR192* as a novel regulator of pancreatic GLP1R. To our knowledge, this observation is the first identification of an intrinsic molecular factor that can contribute to differences in susceptibility to statin-induced diabetes.

While a variety of mechanisms have been proposed to contribute to impaired glucose metabolism and risk for NOD with statin treatment^42,43^, the supporting evidence in humans has been inconclusive. Through electronic health records from the members of KPNC, we identified a well-defined cohort of statin users to assess statin effects on NOD. The NOD susceptible cases had higher fasting glucose levels and non-significantly higher triglyceride levels prior to statin initiation compared to the NOD resistant controls, suggesting greater underlying metabolic dysregulation in patients who were susceptible to developing diabetes on statin. This is consistent with the finding that having elevated baseline glycemic markers is a risk factor for developing statin-induced NOD^3^. The similarity of statin-induced lipid changes between NOD susceptible cases and NOD resistant controls strongly suggests that the risk of NOD was not due to greater statin exposure but rather to inherently heightened susceptibility to developing diabetes that was exacerbated by statin use.

We observed that elevated miR-192 in β-cell models enhanced GLP-1 mediated GSIS, which is counterintuitive to the reports in which higher circulating miR-192 levels have been associated with type 1 and type 2 diabetes^29–32^. However, it remains unclear whether elevated serum miR-192 leads to diabetes or, as our results implied, acts as a compensatory mechanism in response to impaired glycemic control to boost insulin secretion. The only other study that examined the effects of miR-192 in the pancreas concluded that miR-192 suppresses β-cell proliferation and inhibits insulin secretion using an autoimmune type 1 diabetes model of mouse pancreatic β-cell line, NIT-1 ^34^. The discrepancy with our results regarding the effects of miR-192 on GSIS may be attributed to the use of an atypical model, as we demonstrated that miR-192 increased GLP-1 potentiated GSIS in multiple *in vitro* and *ex vivo* rodent β-cell models.

Glycemic control following a meal relies on the secretion of GLP-1 and GIP, incretin hormones released by the intestine. Upon binding to their respective receptors, GLP1R and GIPR, in pancreatic β-cells, these hormones enhance GSIS^44^. Under normal conditions, GIP is the primary hormone facilitating GSIS^45^. However, in the diabetic state, GIPR becomes insensitive, while GLP1R remains functional^46^. Two classes of agents commonly used to treat T2DM function through activation of GLP1R. First, GLP1R agonists (e.g. semaglutide, liraglutide), are GLP-1 mimics that induce a more robust increase in GSIS than GLP-1, which is subject to rapid enzymatic decay by dipeptidyl-peptidase-IV (DPP-IV). Importantly, GLP1R agonists have been shown to not only reduce blood glucose, but also to reduce obesity^47^ and to protect against atherosclerotic cardiovascular disease^48^, the leading cause of death in patients with T2DM^49^. Second, DPP-IV inhibitors (e.g. sitagliptin, vildagliptin) reduce degradation of GLP-1, and therefore their clinical benefit is also mediated by GLP1R^50^. The discovery of novel regulators of GLP1R could be used to inform the development of new therapeutics designed to enhance the effects of GLP1R agonists and reduce the risk of developing common gastrointestinal adverse effects associated with these drugs^51^ by enabling lower effective doses.

Interestingly, although we didn’t observe a direct effect of miR-192 on *Gipr* transcript levels in INS-1 cells, *ex vivo* perifusion of islets from miR-192 AAV treated mice secreted more insulin upon stimulation of GIP in GSIS. This GIP response could be due to overall improvements in β-cell function and survival mediated by GLP1R activation, as previously suggested^52^.

The molecular basis of the GLP1R signaling cascade has been extensively studied^53^, however, considerably less is known regarding the regulation of GLP1R levels. At the protein level, GLP1R is known to undergo rapid homologous desensitization in response to agonist stimulation^54^, while N-glycosylation enhances receptor stability and prolongs its half-life^55^. *In vitro* studies have found that *Glp1r* transcript levels are modulated by high dose (but not low dose) glucose^56^, androgens^57^, and agents that increase cAMP, including GLP-1^54^. In addition, recently miR-204-5p, a highly β-cell enriched miRNA, has been shown to directly target the 3’ UTR of *GLP1R* and downregulate its expression in INS-1 cells and primary mouse and human islets^37^. Importantly, *in vivo* deletion of *Mir204* enhanced responsiveness to GLP1R agonists, resulting in improved glucose tolerance, insulin secretion, and protection against diabetes. These findings demonstrate that GLP1R modifiers can impact key physiological processes in the maintenance of glucose homeostasis *in vivo*. Identifying molecular regulators of GLP1R expression may yield important new insight into factors underlying β-cell dysfunction and variation in response to GLP1R agonists and may inform the development of novel therapeutics to elevate GLP1R agonist efficacy.

The effect of miR-192 on GLP1R was at least in part mediated via the interaction with the *GLP1R* 3’ UTR, as indicated by our luciferase reporter assays. This interaction could involve direct binding of miR-192 to the *GLP1R* 3’ UTR or an indirect effect through miR-192 regulation of, or competition with, another miRNA. The upregulation of GLP1R by miR-192 occurred through a different regulatory site than the one previously reported in the human renal tubular epithelial cell line HK-2^33^ and the human enteroendocrine cell line NCI-H716^34^, indicating tissue specificity. The exact sequence within the GLP1R 3’ UTR that mediates the miR-192 effect, as well as the mechanism by which miR-192 upregulates GLP1R, requires further investigation. The most well-understood mechanism by which miRNAs regulate gene expression is through their partnering with members of the argonaute (AGO) protein family and other associated proteins, like TNRC6 (aka GW182), to form a miRNA-ribonucleoprotein complex (microRNP), known as RNA-induced silencing complex (RISC), and cause gene repression^58^. Instances of miRNAs upregulating gene expression are relatively scarce in the literature, although not entirely absent, and there are several proposed mechanisms underlying this phenomenon. First, miRNAs have been shown to interact with the 5’ UTR of their target mRNA and promote mRNA association with ribosomal proteins resulting in upregulated protein levels^59,60^. Second, miRNAs could compete with factors mediating mRNA degradation or translational inhibition which leads to the loss of repression of gene expression^61,62^. Finally, depending on its components, microRNP can also facilitate translation activation. For instance, it has been shown that MIR369 recruits the activating microRNP consisting of AGO2 and fragile X mentalretardation–related protein 1 (FXR1), leading to the up-regulation of TNFalpha translation^63^.

In summary, we have identified a novel role for *MIR192* in statin-induced NOD through its regulation of β-cell GLP1R and GLP-1 mediated GSIS. This mechanism may have broader relevance to the pathobiology of T2DM and could inform new therapeutic strategies, such as developing agents to increase miR-192 levels in pancreatic β-cells to improve disease management, given the pivotal role of GLP1R activation in the effectiveness of GLP1R agonists. More generally, this study demonstrates the power of transcriptomic analysis in iPSCs derived from well-defined case and control patient cohorts to identify causative mechanisms for individual variation in the effects of drug treatment.

## ACKNOWLEDGMENTS

This work would not have been possible without the contributions of the POST study participants. The authors thank Drs. Naohiro Terada and Katherine Santostefano from the Center for Cellular Reprogramming at the University of Florida for their assistance with the reprogramming and validation of the iPSC lines. We also acknowledge the UCSF MLK Cores Research Facility for sample analysis, and the Northwest Genomics Center for generating the iPSC RNA-seq data. This project was supported by the National Institutes of Health Pharmacogenomics Research Network (PGRN) [P50 GM115318], the Canadian Islet Research and Training Network (CIRTN) NSERC CREATE program, the Ontario Graduate Scholarship, and the Diabetes Canada End Diabetes Award, funded by the River Philip Foundation. The funders had no role in the study design, data analysis, or manuscript preparation.

## AUTHOR CONTRIBUTIONS

Conceptualization – RMK, EEM, MWM

Methodology – YK, CAAL, MN, AOO, MWM

Formal Analysis – YK, CAAL, YQ, YZ, ET, ML

Investigation – YK, CAAL, YQ, YZ, ET, AMH, MN, TY, XW, GN,

Resources – GS, ANM, CI

Data Curation – ET, ML

Writing – Original Draft – YK, MWM

Writing – Review & Editing – all authors

Visualization – YK, CAAL

Supervision – CI, EEM, MWM

Project administration - MWM

Funding acquisition – CI, RMK, EEM, MWM

## DECLARATION OF INTERESTS

The authors declare no competing interests.

## Notes

### Competing Interest Statement

The authors have declared no competing interest.

## REFERENCES

1. Cholesterol Treatment Trialists’ (Ctt) Collaborators (2012). The effects of lowering LDL cholesterol with statin therapy in people at low risk of vascular disease: meta-analysis of individual data from 27 randomised trials. The Lancet 380, 581–590. 10.1016/S0140-6736(12)60367-5.

2. Bredefeld, C.L., Choi, P., Cullen, T., Nicolich-Henkin, S.J., and Waters, L. (2024). Statin Use and Hyperglycemia: Do Statins Cause Diabetes? Curr. Atheroscler. Rep. 27, 18. 10.1007/s11883-024-01266-8.

3. Cholesterol Treatment Trialists’ (CTT) Collaboration. Electronic address: and Cholesterol Treatment Trialists’ (CTT) Collaboration (2024). Effects of statin therapy on diagnoses of new-onset diabetes and worsening glycaemia in large-scale randomised blinded statin trials: an individual participant data meta-analysis. Lancet Diabetes Endocrinol. 12, 306–319. 10.1016/S2213-8587(24)00040-8.

4. Grundy, S.M., Stone, N.J., Bailey, A.L., Beam, C., Birtcher, K.K., Blumenthal, R.S., Braun, L.T., De Ferranti, S., Faiella-Tommasino, J., Forman, D.E., et al. (2019). 2018 AHA/ACC/AACVPR/AAPA/ABC/ACPM/ADA/AGS/APhA/ASPC/NLA/PCNA Guideline on the Management of Blood Cholesterol: Executive Summary. J. Am. Coll. Cardiol. 73, 3168–3209. 10.1016/j.jacc.2018.11.002.

5. Cosentino, F., Grant, P.J., Aboyans, V., Bailey, C.J., Ceriello, A., Delgado, V., Federici, M., Filippatos, G., Grobbee, D.E., Hansen, T.B., et al. (2020). 2019 ESC Guidelines on diabetes, pre-diabetes, and cardiovascular diseases developed in collaboration with the EASD. Eur. Heart J. 41, 255–323. 10.1093/eurheartj/ehz486.

6. Rosoff, D.B., Wagner, J., Jung, J., Pacher, P., Christodoulides, C., Smith, G.D., Ray, D., and Lohoff, F.W. (2024). Multi-Omics Mendelian Randomization Study Investigating the Impact of PCSK9 and HMGCR Inhibition on Type 2 Diabetes Across Five Populations. Diabetes, db240451. 10.2337/db24-0451.

7. She, J., Tuerhongjiang, G., Guo, M., Liu, J., Hao, X., Guo, L., Liu, N., Xi, W., Zheng, T., Du, B., et al. (2024). Statins aggravate insulin resistance through reduced blood glucagon-like peptide-1 levels in a microbiota-dependent manner. Cell Metab. 36, 408-421.e5. 10.1016/j.cmet.2023.12.027.

8. Brubaker, P.L., and Drucker, D.J. (2004). Minireview: Glucagon-Like Peptides Regulate Cell Proliferation and Apoptosis in the Pancreas, Gut, and Central Nervous System. Endocrinology 145, 2653–2659. 10.1210/en.2004-0015.

9. Baggio, L.L., and Drucker, D.J. (2007). Biology of Incretins: GLP-1 and GIP. Gastroenterology 132, 2131–2157. 10.1053/j.gastro.2007.03.054.

10. Li, Y., and Rosenblit, P.D. (2018). Glucagon-Like Peptide-1 Receptor Agonists and Cardiovascular Risk Reduction in Type 2 Diabetes Mellitus: Is It a Class Effect? Curr. Cardiol. Rep. 20, 113. 10.1007/s11886-018-1051-2.

11. Hinnen, D. (2017). Glucagon-Like Peptide 1 Receptor Agonists for Type 2 Diabetes. Diabetes Spectr. 30, 202–210. 10.2337/ds16-0026.

12. Popoviciu, M.-S., Păduraru, L., Yahya, G., Metwally, K., and Cavalu, S. (2023). Emerging Role of GLP-1 Agonists in Obesity: A Comprehensive Review of Randomised Controlled Trials. Int. J. Mol. Sci. 24, 10449. 10.3390/ijms241310449.

13. Lincoff, A.M., Brown-Frandsen, K., Colhoun, H.M., Deanfield, J., Emerson, S.S., Esbjerg, S., Hardt-Lindberg, S., Hovingh, G.K., Kahn, S.E., Kushner, R.F., et al. (2023). Semaglutide and Cardiovascular Outcomes in Obesity without Diabetes. N. Engl. J. Med. 389, 2221–2232. 10.1056/NEJMoa2307563.

14. Scattolini, V., Luni, C., Zambon, A., Galvanin, S., Gagliano, O., Ciubotaru, C.D., Avogaro, A., Mammano, F., Elvassore, N., and Fadini, G.P. (2016). Simvastatin Rapidly and Reversibly Inhibits Insulin Secretion in Intact Single-Islet Cultures. Diabetes Ther. 7, 679–693. 10.1007/s13300-016-0210-y.

15. Urbano, F., Bugliani, M., Filippello, A., Scamporrino, A., Di Mauro, S., Di Pino, A., Scicali, R., Noto, D., Rabuazzo, A.M., Averna, M., et al. (2017). Atorvastatin but Not Pravastatin Impairs Mitochondrial Function in Human Pancreatic Islets and Rat β-Cells. Direct Effect of Oxidative Stress. Sci. Rep. 7, 11863. 10.1038/s41598-017-11070-x.

16. Yaluri, N., Modi, S., López Rodríguez, M., Stančáková, A., Kuusisto, J., Kokkola, T., and Laakso, M. (2015). Simvastatin Impairs Insulin Secretion by Multiple Mechanisms in MIN6 Cells. PLOS ONE 10, e0142902. 10.1371/journal.pone.0142902.

17. Avior, Y., Sagi, I., and Benvenisty, N. (2016). Pluripotent stem cells in disease modelling and drug discovery. Nat. Rev. Mol. Cell Biol. 17, 170–182. 10.1038/nrm.2015.27.

18. Sterneckert, J.L., Reinhardt, P., and Schöler, H.R. (2014). Investigating human disease using stem cell models. Nat. Rev. Genet. 15, 625–639. 10.1038/nrg3764.

19. Kuang, Y.-L., Munoz, A., Nalula, G., Santostefano, K.E., Sanghez, V., Sanchez, G., Terada, N., Mattis, A.N., Iacovino, M., Iribarren, C., et al. (2019). Evaluation of commonly used ectoderm markers in iPSC trilineage differentiation. Stem Cell Res. 37, 101434. 10.1016/j.scr.2019.101434.

20. Okita, K., Yamakawa, T., Matsumura, Y., Sato, Y., Amano, N., Watanabe, A., Goshima, N., and Yamanaka, S. (2013). An Efficient Nonviral Method to Generate Integration-Free Human-Induced Pluripotent Stem Cells from Cord Blood and Peripheral Blood Cells. Stem Cells 31, 458–466. 10.1002/stem.1293.

21. Fadzeyeva, E., Locatelli, C.A.A., Trzaskalski, N.A., Nguyen, M.-A., Capozzi, M.E., Vulesevic, B., Morrow, N.M., Ghorbani, P., Hanson, A.A., Lorenzen-Schmidt, I., et al. (2023). Pancreas-derived DPP4 is not essential for glucose homeostasis under metabolic stress. iScience 26, 106748. 10.1016/j.isci.2023.106748.

22. Dobin, A., Davis, C.A., Schlesinger, F., Drenkow, J., Zaleski, C., Jha, S., Batut, P., Chaisson, M., and Gingeras, T.R. (2013). STAR: ultrafast universal RNA-seq aligner. Bioinformatics 29, 15–21. 10.1093/bioinformatics/bts635.

23. Liao, Y., Smyth, G.K., and Shi, W. (2014). featureCounts: an efficient general purpose program for assigning sequence reads to genomic features. Bioinformatics 30, 923–930. 10.1093/bioinformatics/btt656.

24. Love, M.I., Huber, W., and Anders, S. (2014). Moderated estimation of fold change and dispersion for RNA-seq data with DESeq2. Genome Biol. 15, 550. 10.1186/s13059-014-0550-8.

25. Hansotia, T., Baggio, L.L., Delmeire, D., Hinke, S.A., Yamada, Y., Tsukiyama, K., Seino, Y., Holst, J.J., Schuit, F., and Drucker, D.J. (2004). Double Incretin Receptor Knockout (DIRKO) Mice Reveal an Essential Role for the Enteroinsular Axis in Transducing the Glucoregulatory Actions of DPP-IV Inhibitors. Diabetes 53, 1326–1335. 10.2337/diabetes.53.5.1326.

26. Dong, B., Wu, M., Cao, A., Li, H., and Liu, J. (2010). Suppression of Idol expression is an additional mechanism underlying statin-induced up-regulation of hepatic LDL receptor expression. Int. J. Mol. Med. 10.3892/ijmm.2010.559.

27. Goldstein, J.L., and Brown, M.S. (1990). Regulation of the mevalonate pathway. Nature 343, 425–430. 10.1038/343425a0.

28. Oni-Orisan, A., Hoffmann, T.J., Ranatunga, D., Medina, M.W., Jorgenson, E., Schaefer, C., Krauss, R.M., Iribarren, C., and Risch, N. (2018). Characterization of Statin Low-Density Lipoprotein Cholesterol Dose-Response Using Electronic Health Records in a Large Population-Based Cohort. Circ. Genomic Precis. Med. 11, e002043. 10.1161/CIRCGEN.117.002043.

29. Párrizas, M., Brugnara, L., Esteban, Y., González-Franquesa, A., Canivell, S., Murillo, S., Gordillo-Bastidas, E., Cussó, R., Cadefau, J.A., García-Roves, P.M., et al. (2015). Circulating miR-192 and miR-193b Are Markers of Prediabetes and Are Modulated by an Exercise Intervention. J. Clin. Endocrinol. Metab. 100, E407–E415. 10.1210/jc.2014-2574.

30. Jaeger, A., Zollinger, L., Saely, C.H., Muendlein, A., Evangelakos, I., Nasias, D., Charizopoulou, N., Schofield, J.D., Othman, A., Soran, H., et al. (2018). Circulating microRNAs -192 and -194 are associated with the presence and incidence of diabetes mellitus. Sci. Rep. 8, 14274. 10.1038/s41598-018-32274-9.

31. Erener, S., Marwaha, A., Tan, R., Panagiotopoulos, C., and Kieffer, T.J. (2017). Profiling of circulating microRNAs in children with recent onset of type 1 diabetes. JCI Insight 2. 10.1172/jci.insight.89656.

32. Yang, Z., Chen, H., Si, H., Li, X., Ding, X., Sheng, Q., Chen, P., and Zhang, H. (2014). Serum miR-23a, a potential biomarker for diagnosis of pre-diabetes and type 2 diabetes. Acta Diabetol. 51, 823–831. 10.1007/s00592-014-0617-8.

33. Jia, Y., Zheng, Z., Guan, M., Zhang, Q., Li, Y., Wang, L., and Xue, Y. (2018). Exendin-4 ameliorates high glucose-induced fibrosis by inhibiting the secretion of miR-192 from injured renal tubular epithelial cells. Exp. Mol. Med. 50, 1–13. 10.1038/s12276-018-0084-3.

34. Pan, W., Zhang, Y., Zeng, C., Xu, F., Yan, J., and Weng, J. (2018). miR-192 is upregulated in T1DM, regulates pancreatic β-cell development and inhibits insulin secretion through suppressing GLP-1 expression. Exp. Ther. Med. 10.3892/etm.2018.6453.

35. Ludwig, N., Leidinger, P., Becker, K., Backes, C., Fehlmann, T., Pallasch, C., Rheinheimer, S., Meder, B., Stähler, C., Meese, E., et al. (2016). Distribution of miRNA expression across human tissues. Nucleic Acids Res. 44, 3865–3877. 10.1093/nar/gkw116.

36. Van De Bunt, M., Gaulton, K.J., Parts, L., Moran, I., Johnson, P.R., Lindgren, C.M., Ferrer, J., Gloyn, A.L., and McCarthy, M.I. (2013). The miRNA Profile of Human Pancreatic Islets and Beta-Cells and Relationship to Type 2 Diabetes Pathogenesis. PLoS ONE 8, e55272. 10.1371/journal.pone.0055272.

37. Jo, S., Chen, J., Xu, G., Grayson, T.B., Thielen, L.A., and Shalev, A. (2018). miR-204 Controls Glucagon-Like Peptide 1 Receptor Expression and Agonist Function. Diabetes 67, 256–264. 10.2337/db17-0506.

38. Ast, J., Broichhagen, J., and Hodson, D.J. (2021). Reagents and models for detecting endogenous GLP1R and GIPR. EBioMedicine 74, 103739. 10.1016/j.ebiom.2021.103739.

39. Bjørnholm, K.D., Povlsen, G.K., Ougaard, M.E., Pyke, C., Rakipovski, G., Tveden-Nyborg, P., Lykkesfeldt, J., and Skovsted, G.F. (2021). Decreased expression of the GLP-1 receptor after segmental artery injury in mice. J. Endocrinol. 248, 289–301. 10.1530/JOE-20-0608.

40. Pauza, A.G., Thakkar, P., Tasic, T., Felippe, I., Bishop, P., Greenwood, M.P., Rysevaite-Kyguoliene, K., Ast, J., Broichhagen, J., Hodson, D.J., et al. (2022). GLP1R Attenuates Sympathetic Response to High Glucose via Carotid Body Inhibition. Circ. Res. 130, 694–707. 10.1161/CIRCRESAHA.121.319874.

41. Liu, X.-L., Cao, H.-X., Wang, B.-C., Xin, F.-Z., Zhang, R.-N., Zhou, D., Yang, R.-X., Zhao, Z.-H., Pan, Q., and Fan, J.-G. (2017). miR-192-5p regulates lipid synthesis in non-alcoholic fatty liver disease through SCD-1. World J. Gastroenterol. 23, 8140–8151. 10.3748/wjg.v23.i46.8140.

42. Guber, K., Pemmasani, G., Malik, A., Aronow, W.S., Yandrapalli, S., and Frishman, W.H. (2021). Statins and Higher Diabetes Mellitus Risk: Incidence, Proposed Mechanisms, and Clinical Implications. Cardiol. Rev. 29, 314–322. 10.1097/CRD.0000000000000348.

43. Yandrapalli, S., Malik, A., Guber, K., Rochlani, Y., Pemmasani, G., Jasti, M., and Aronow, W.S. (2019). Statins and the potential for higher diabetes mellitus risk. Expert Rev. Clin. Pharmacol. 12, 825–830. 10.1080/17512433.2019.1659133.

44. Komatsu, M., Takei, M., Ishii, H., and Sato, Y. (2013). Glucose-stimulated insulin secretion: A newer perspective. J. Diabetes Investig. 4, 511–516. 10.1111/jdi.12094.

45. Nauck, M.A., Bartels, E., Orskov, C., Ebert, R., and Creutzfeldt, W. (1993). Additive insulinotropic effects of exogenous synthetic human gastric inhibitory polypeptide and glucagon-like peptide-1-(7-36) amide infused at near-physiological insulinotropic hormone and glucose concentrations. J. Clin. Endocrinol. Metab. 76, 912–917. 10.1210/jcem.76.4.8473405.

46. Nauck, M.A., Heimesaat, M.M., Orskov, C., Holst, J.J., Ebert, R., and Creutzfeldt, W. (1993). Preserved incretin activity of glucagon-like peptide 1 [7-36 amide] but not of synthetic human gastric inhibitory polypeptide in patients with type-2 diabetes mellitus. J. Clin. Invest. 91, 301–307. 10.1172/JCI116186.

47. Van Gaal, L., and Scheen, A. (2015). Weight Management in Type 2 Diabetes: Current and Emerging Approaches to Treatment. Diabetes Care 38, 1161–1172. 10.2337/dc14-1630.

48. Lim, G.B. (2019). GLP1R agonists: primary cardiovascular prevention and oral administration. Nat. Rev. Cardiol. 16, 453–453. 10.1038/s41569-019-0232-z.

49. Einarson, T.R., Acs, A., Ludwig, C., and Panton, U.H. (2018). Prevalence of cardiovascular disease in type 2 diabetes: a systematic literature review of scientific evidence from across the world in 2007– 2017. Cardiovasc. Diabetol. 17, 83. 10.1186/s12933-018-0728-6.

50. Gilbert, M.P., and Pratley, R.E. (2020). GLP-1 Analogs and DPP-4 Inhibitors in Type 2 Diabetes Therapy: Review of Head-to-Head Clinical Trials. Front. Endocrinol. 11, 178. 10.3389/fendo.2020.00178.

51. Bettge, K., Kahle, M., Abd El Aziz, M.S., Meier, J.J., and Nauck, M.A. (2017). Occurrence of nausea, vomiting and diarrhoea reported as adverse events in clinical trials studying glucagon-like peptide-1 receptor agonists: A systematic analysis of published clinical trials. Diabetes Obes. Metab. 19, 336–347. 10.1111/dom.12824.

52. Yusta, B., Baggio, L.L., Estall, J.L., Koehler, J.A., Holland, D.P., Li, H., Pipeleers, D., Ling, Z., and Drucker, D.J. (2006). GLP-1 receptor activation improves beta cell function and survival following induction of endoplasmic reticulum stress. Cell Metab. 4, 391–406. 10.1016/j.cmet.2006.10.001.

53. Nadkarni, P., Chepurny, O.G., and Holz, G.G. (2014). Regulation of Glucose Homeostasis by GLP-1. In Progress in Molecular Biology and Translational Science (Elsevier), pp. 23–65. 10.1016/B978-0-12-800101-1.00002-8.

54. Fehmann, H.-C., Jiang, J., Pitt, D., Schweinfurth, J., and Göke, B. (1996). Ligand-Induced Regulation of Glucagon-like Peptide-I Receptor Function and Expression in Insulin-Secreting β Cells: Pancreas 13, 273–282. 10.1097/00006676-199610000-00010.

55. Whitaker, G.M., Lynn, F.C., McIntosh, C.H.S., and Accili, E.A. (2012). Regulation of GIP and GLP1 Receptor Cell Surface Expression by N-Glycosylation and Receptor Heteromerization. PLoS ONE 7, e32675. 10.1371/journal.pone.0032675.

56. Abrahamsen, N., and Nishimura, E. (1995). Regulation of glucagon and glucagon-like peptide-1 receptor messenger ribonucleic acid expression in cultured rat pancreatic islets by glucose, cyclic adenosine 3’,5’-monophosphate, and glucocorticoids. Endocrinology 136, 1572–1578. 10.1210/endo.136.4.7534705.

57. Zhu, L., Zhou, J., Pan, Y., Lv, J., Liu, Y., Yu, S., and Zhang, Y. (2019). Glucagon-like peptide-1 receptor expression and its functions are regulated by androgen. Biomed. Pharmacother. 120, 109555. 10.1016/j.biopha.2019.109555.

58. Iwakawa, H., and Tomari, Y. (2022). Life of RISC: Formation, action, and degradation of RNA-induced silencing complex. Mol. Cell 82, 30–43. 10.1016/j.molcel.2021.11.026.

59. Ørom, U.A., Nielsen, F.C., and Lund, A.H. (2008). MicroRNA-10a Binds the 5′UTR of Ribosomal Protein mRNAs and Enhances Their Translation. Mol. Cell 30, 460–471. 10.1016/j.molcel.2008.05.001.

60. Tsai, N.-P., Lin, Y.-L., and Wei, L.-N. (2009). MicroRNA mir-346 targets the 5′-untranslated region of receptor-interacting protein 140 (RIP140) mRNA and up-regulates its protein expression. Biochem. J. 424, 411–418. 10.1042/BJ20090915.

61. Murphy, A.J., Guyre, P.M., and Pioli, P.A. (2010). Estradiol Suppresses NF-κB Activation through Coordinated Regulation of let-7a and miR-125b in Primary Human Macrophages. J. Immunol. 184, 5029–5037. 10.4049/jimmunol.0903463.

62. Ma, F., Liu, X., Li, D., Wang, P., Li, N., Lu, L., and Cao, X. (2010). MicroRNA-466l Upregulates IL-10 Expression in TLR-Triggered Macrophages by Antagonizing RNA-Binding Protein Tristetraprolin-Mediated IL-10 mRNA Degradation. J. Immunol. 184, 6053–6059. 10.4049/jimmunol.0902308.

63. Vasudevan, S., Tong, Y., and Steitz, J.A. (2007). Switching from Repression to Activation: MicroRNAs Can Up-Regulate Translation. Science 318, 1931–1934. 10.1126/science.1149460.

