## Supplementary Figures for "*MIR192* Upregulates GLP-1 Receptor and Improves Statin-Induced Impairment of Insulin Secretion"

**A** NOD Susceptible Cases

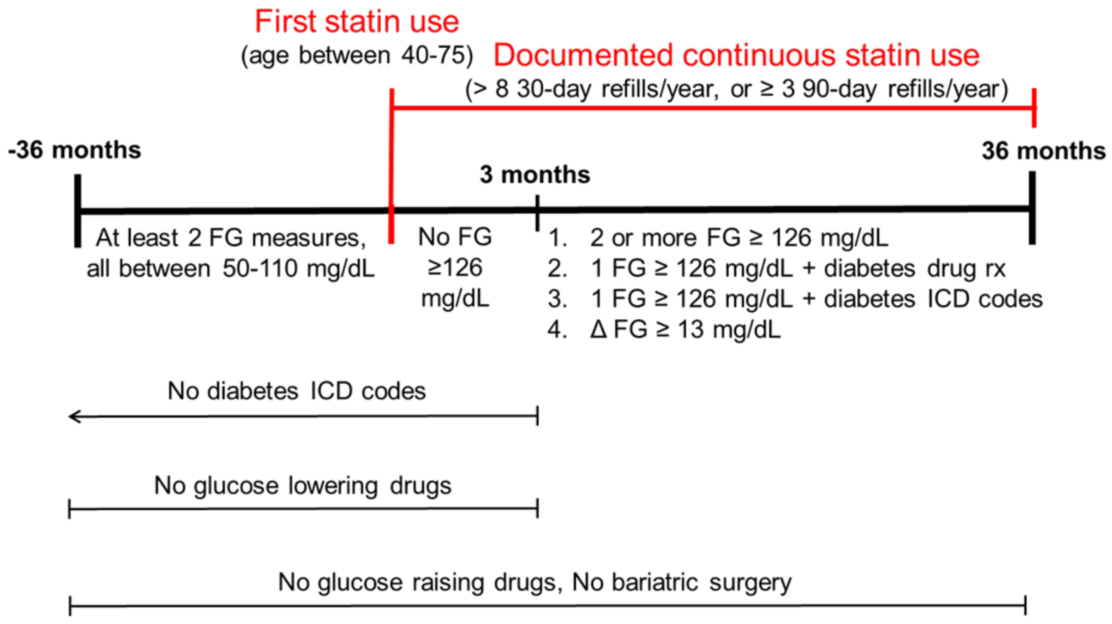


**B** NOD Resistant Controls

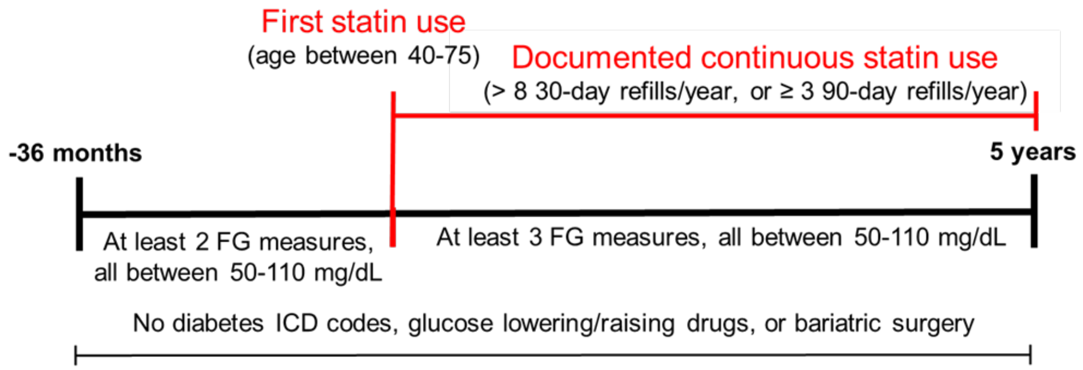


**Supplementary Figure S1. Algorithm used to identify stain-induced new-onset diabetes (NOD) susceptible case (A) or NOD resistant control (B) cohorts from the electronic databases of the Kaiser Permanente of Northern California.** Abbreviations: FG, fasting glucose; ICD, international classification of diseases; rx, prescription.

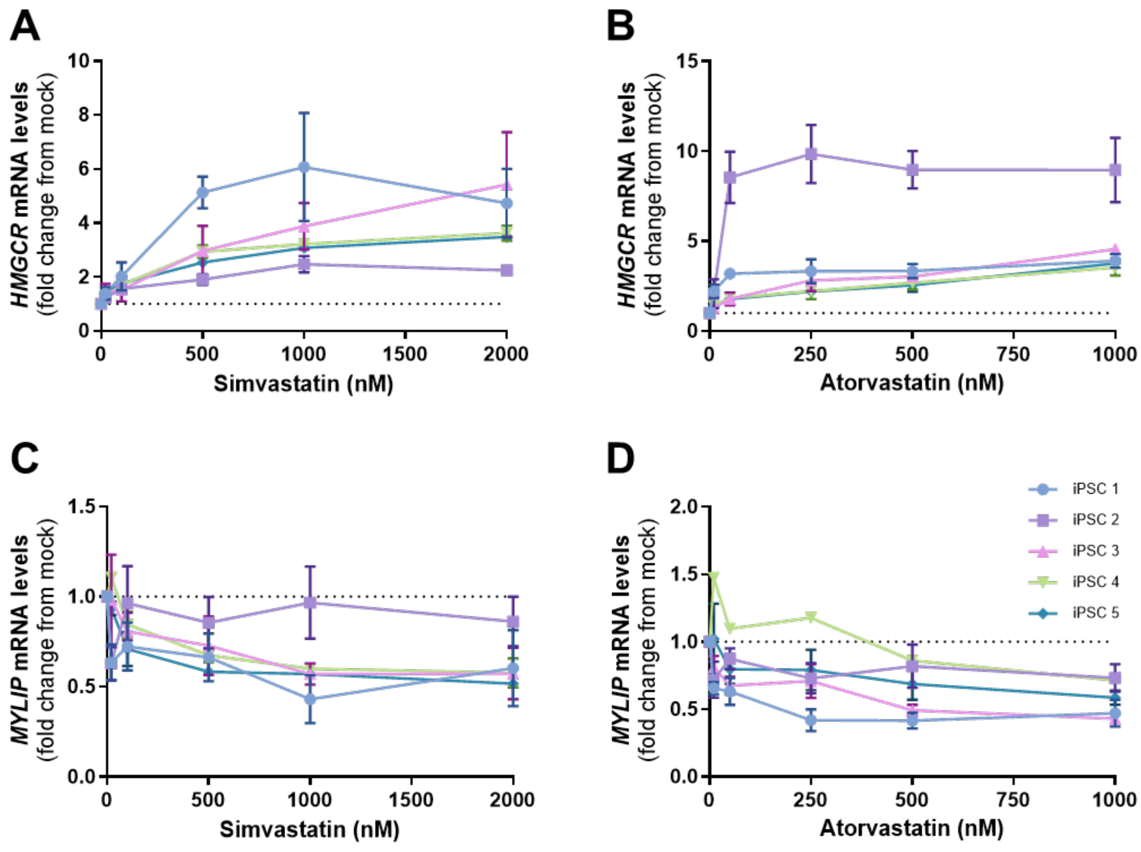


**Supplementary Figure S2. Statin dose response on *HMGCR* and *MYLIP* expression in iPSC.** Five iPSC lines (n=1-4) were treated with a wide concentration range of simvastatin and atorvastatin for 24 hours before *HMGCR* (**A-B**) and *MYLIP* (**C-D**) transcript levels were quantified by qPCR and normalized to *CLPTM1*. Data are presented as mean +/- SEM.

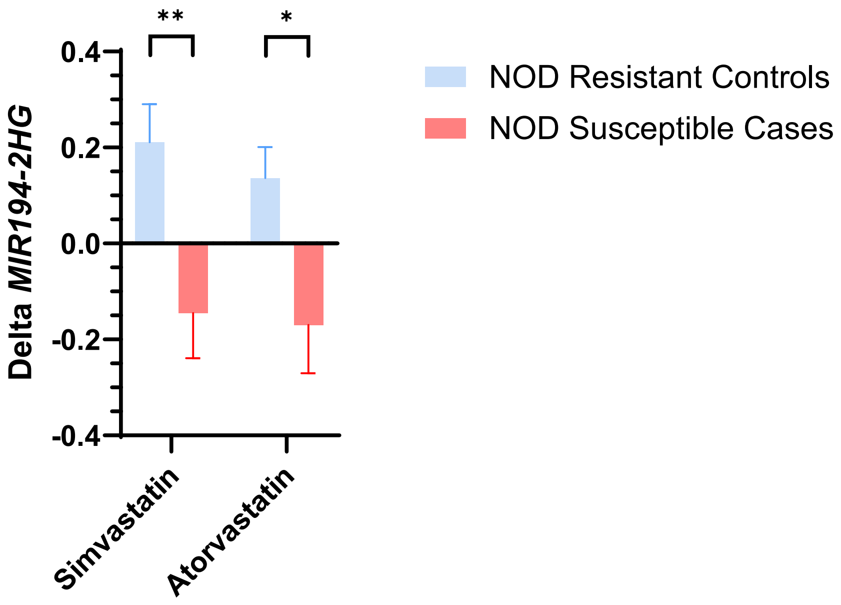


**Supplementary Figure S3**. Change in *MIR194-2HG* levels in iPSCs derived from statin users whose pre-statin fasting glucose levels were less than 100 mg/dL and categorized as NOD susceptible cases (n=19) or NOD resistant controls (n=13) after 24hr incubation with 500 nM simvastatin, 250 nM atorvastatin or mock control buffer. Values were variance stabilized (~log2 transformation) in DESeq2 and deltas were calculated as statin-control expression levels. Data are presented as mean +/- SEM, significance was determined by unpaired t-test, *$p < 0.05$, **$p < 0.01$.

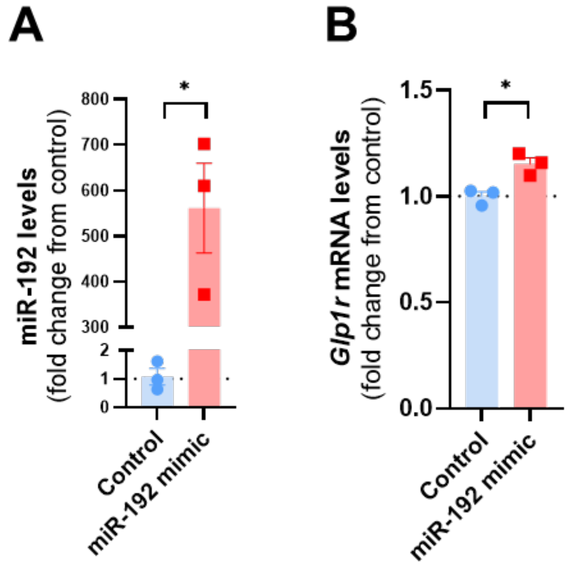


**Supplementary Figure S4**. **miR-192 upregulates *Glp1r* transcript levels in βTC3 cells.** Mouse βTC3 cells were transfected with 10 nM miR-192 mimic or a non-targeting control. After 48 hours, miR-192 (**A**) and *Glp1r* (**B**) transcript levels were quantified by qPCR and normalized to miR-16 and *Clptm1,* respectively. Data are presented as mean +/- SEM, significance was determined by unpaired t-test, *$p < 0.05$.

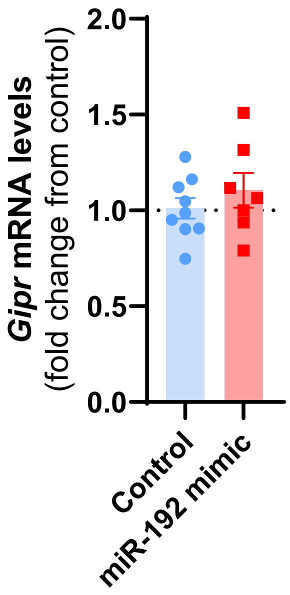


**Supplementary Figure S5**. **miR-192 regulation of *Gipr* in INS-1.** INS-1 rat insulinoma cells were transfected with 10 nM miR-192 mimic or a non-targeting control for 48 hours. *Gipr* transcript levels were quantified by qPCR, normalized to the house keeping gene *Clptm1*. Data are presented as mean +/- SEM.

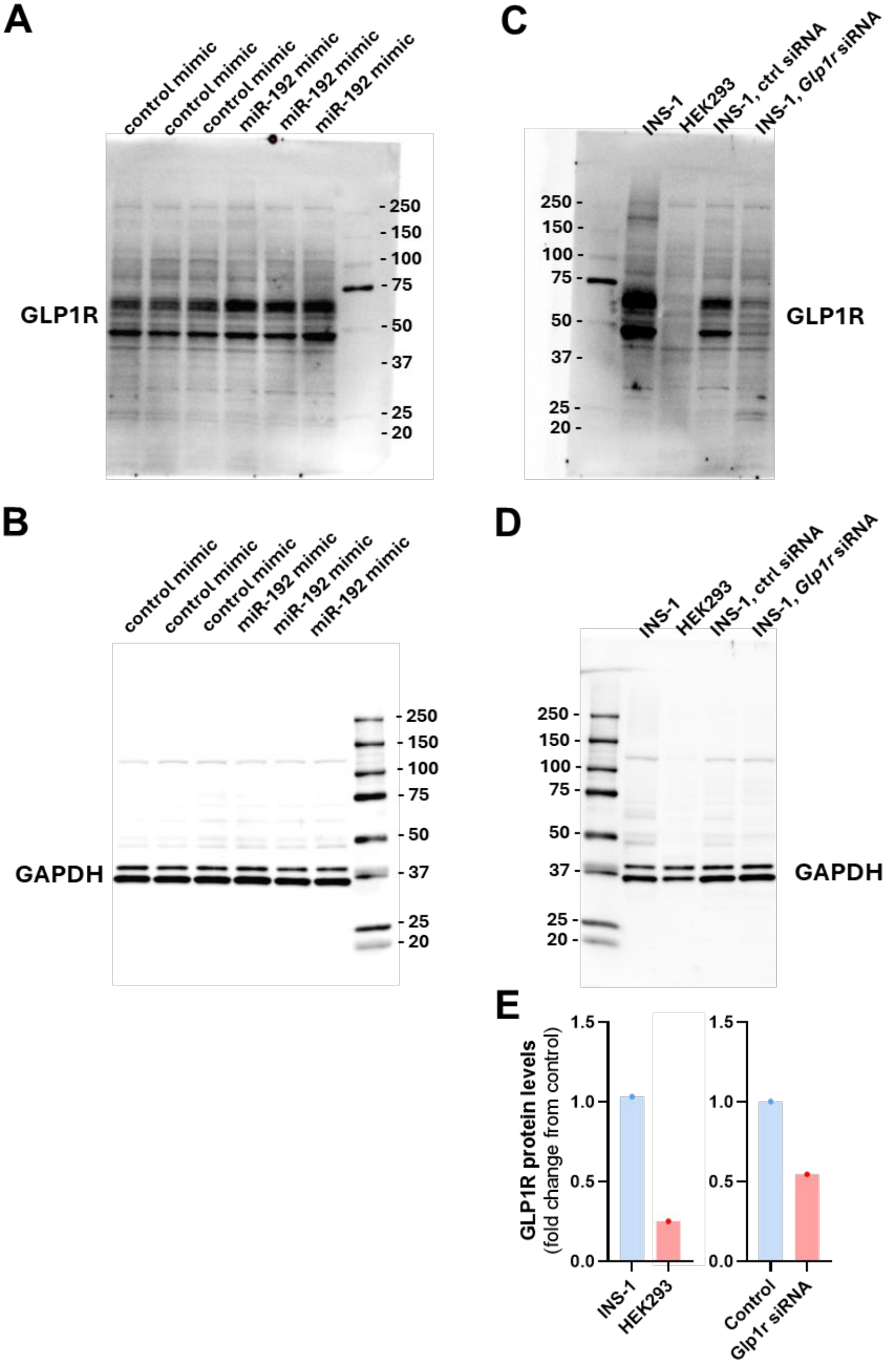


**Supplementary Figure S6. Western blot images and GLP1R antibody validation. (A-B)** Whole blot images of the GLP1R and GAPDH western blots shown in **Figure 2F**. **(C-D)** To test the specificity of the GLP1R antibody for use in western blots, cell lysates were collected from INS-1 and HEK293 cells, or INS-1 cells treated with either Glp1r siRNA or a non-targeting negative control siRNA. (**E**) Bar graphs of the western blot quantification shown in **Fig. S6C-D**. GLP1R protein levels were normalized to GAPDH and expressed as fold change from INS-1 or Control siRNA treated samples.

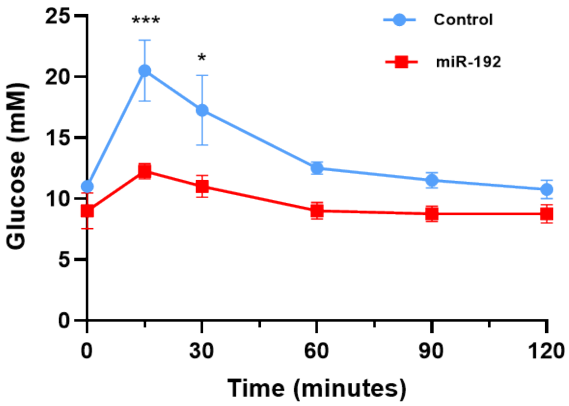


**Supplementary Figure S7.** Seven weeks old wild-type C57BL/6J male mice (n=4/group) were fed a GAN diet (Research Diets, #D09100310, 40 kcal% fat, 20 kcal% fructose and 2% cholesterol) for 4 weeks before either miR-192 or control AAV vectors ($8x10^{12}$ viral genomes/kg) were delivered via intraperitoneal injection. Four more weeks later, intraperitoneal glucose tolerance test was performed with 1.5 g/kg glucose after a 6 hour fast, and plasma glucose was determined over 2 hours. Data are presented as mean +/- SEM, significance was determined by two-way ANOVA with Šídák's correction for multiple comparisons, *$p < 0.05$, ***$p < 0.001$.

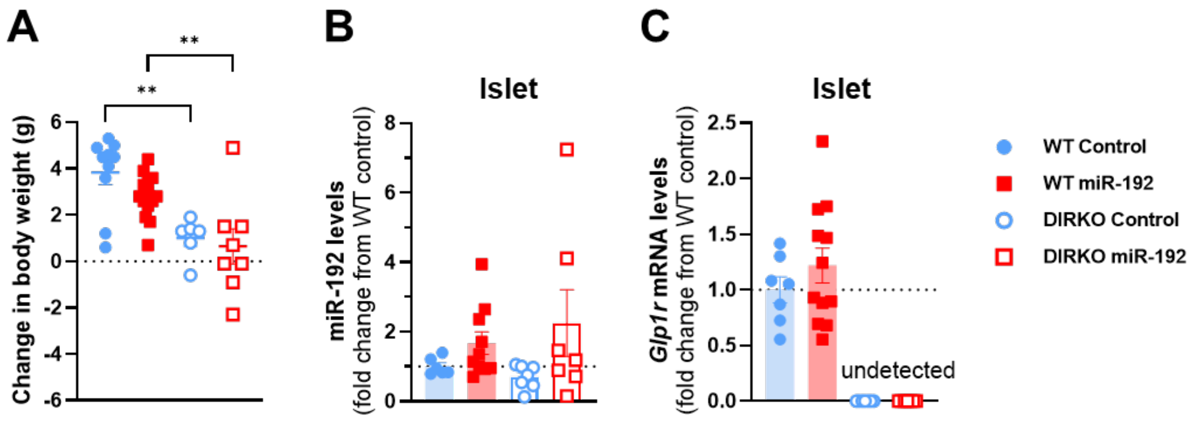


**Supplementary Figure S8.** Male C57BL/6J wild-type and double incretin receptor knockout (DIRKO) mice were placed on a high fat diet (42 kcal% fat and 0.2% cholesterol) at 12 weeks of age. After 8 weeks on this diet, they received a one-time tail vein injection of either miR-192 (AAV8-EF1α-mmu-mir-192-eGFP, Vector Biolabs) or a scrambled control (scAAV8-EF1α-ctrl-miR-eGFP, Vector Biolabs) AAV at a dose of 1 x $10^{11}$ GC per mouse in 200 µl. (**A**) Body mass change between the 4 weeks from tail vein injection to the first metabolic test. (**B-C**) Relative abundance of miR-192 levels normalized to miR-16 (**B**) and *Glp1r* transcript levels normalized to *ActB* (**C**) in islets from wildtype and DIRKO mice. Data are presented as mean +/- SEM, significance was determined by one-way ANOVA with Šídák's correction for multiple comparisons, $**p < 0.01$.

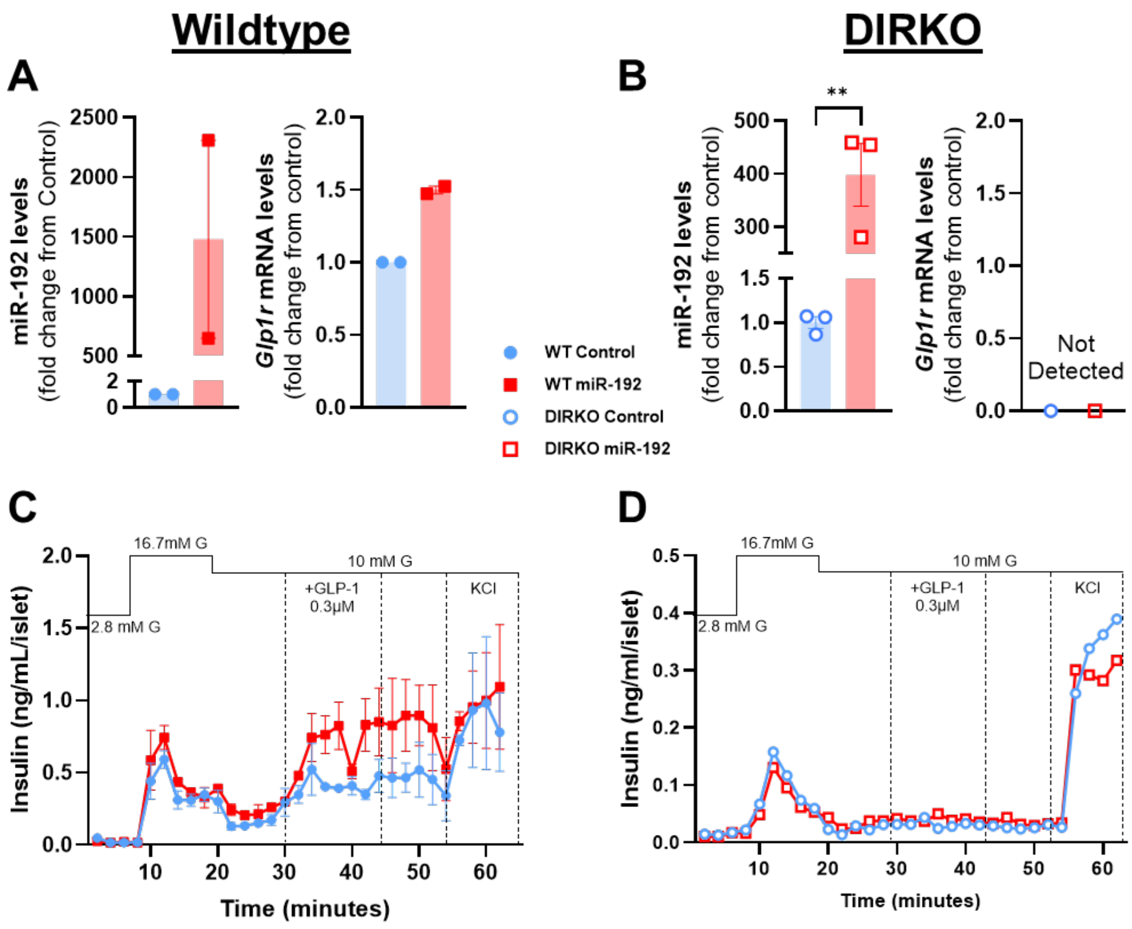


**Supplementary Figure S9. miR-192 mimic increases *Glp1r* transcript levels and GLP-1 augmented glucose-stimulated insulin secretion (GSIS) in wildtype primary mouse islets.** Primary islets from wildtype (WT) mice or double incretin receptor knockout (DIRKO) mice were dispersed, transfected with the miR-192 mimic or control, reaggregated, and **(A-B)** miR-192 and *Glp1r* transcript levels were quantified by qPCR and normalized to miR-16 and *Actb*, respectively. **(C-D)** GSIS was quantified by perifusion, n=2/group for wildtype islets **(C),** and n=1/group for DIRKO islets **(D)**. Significance could not be calculated for wildtype islets due to insufficient number of replicates but was determined for miR-192 levels in DIRKO islets by unpaired t-test, ***p* < 0.01.
